# Lateral Root Priming Synergystically Arises from Root Growth and Auxin Transport Dynamics

**DOI:** 10.1101/361709

**Authors:** Thea van den Berg, Kirsten H. ten Tusscher

## Abstract

The root system is a major determinant of plant fitness. Its capacity to supply the plant with sufficient water and nutrients strongly depends on root system architecture, which arises from the repeated branching off of lateral roots. A critical first step in lateral root formation is priming, which prepatterns sites competent of forming a lateral root. Priming is characterized by temporal oscillations in auxin, auxin signalling and gene expression in the root meristem, which through growth become transformed into a spatially repetitive pattern of competent sites. Previous studies have demonstrated the importance of auxin synthesis, transport and perception for the amplitude of these oscillations and their chances of producing an actual competent site. Additionally, repeated lateral root cap apoptosis was demonstrated to be strongly correlated with repetitive lateral root priming. Intriguingly, no single mutation has been identified that fully abolishes lateral root formation, and thusfar the mechanism underlying oscillations has remained unknown. In this study, we investigated the impact of auxin reflux loop properties combined with root growth dynamics on priming, using a computational approach. To this end we developed a novel multi-scale root model incorporating a realistic root tip architecture and reflux loop properties as well as root growth dynamics. Excitingly, in this model, repetitive auxin elevations automatically emerge. First, we show that root tip architecture and reflux loop properties result in an auxin loading zone at the start of the elongation zone, with preferential auxin loading in narrow vasculature cells. Second, we demonstrate how meristematic root growth dynamics causes regular alternations in the sizes of cells arriving at the elongation zone, which subsequently become amplified during cell expansion. These cell size differences translate into differences in cellular auxin loading potential. Combined, these properties result in temporal and spatial fluctuations in auxin levels in vasculature and pericycle cells. Our model predicts that temporal priming frequency predominantly depends on cell cycle duration, while cell cycle duration together with meristem size control lateral root spacing.

## 1 Introduction

Roots provide plants with anchorage to their substrate, as well as access to water and nutrients. Overall root system architecture (RSA) is thus a major determinant of plant fitness [Herder et al., 2010, Eshel, A., Beeckman, 2013, Kong et al., 2014, Kochian, 2016]. RSA depends both on main root (MR) length as well as the location, number, angle and length of lateral and adventitious roots [Rogers and Benfey, 2015]. RSA is an extremely plastic trait and enables plant survival in a variable environment [Eshel, A., Beeckman, 2013, Rogers and Benfey, 2015]. Nonetheless, major parts of the basic developmental programs on which environmental factors impinge remain poorly understood. In plants with a tap root system, such as most trees, barley and *Arabidopsis*, lateral roots (LRs) emerging from the MR are the major source for RSA branching. The outgrowth of a LR from the MR is preceded by a sequence of developmental processes, starting with the priming of competent cells capable of future LR formation and ending with the emergence of the LR from inside the MR [Malamy and Benfey, 1997, Dubrovsky et al., 2008, Bielach and Benkova, De Smet et al., 2010, De Rybel et al., 2010, Goh et al., 2012]. Here, we focus on this earliest step of LR development, priming.

The mechanisms underlying LR priming have been the subject of intensive investigations in the model plant species *Arabidopsis.* So far, no single mutations have been identified that completely abolish LR formation, indicating that LR formation has a highly robust nature. Instead, the strongest repression of LR formation occurs in the dark, known to affect sugar transport and consequently root growth dynamics [Jensen et al., 1998]. Intriguingly, various investigations have pointed to semi-regular temporal oscillations in the plant hormone auxin and/or its downstream responses as well as gene expression levels to temporally prepattern priming sites [De Smet et al., 2007, Xuan et al., 2015, Moreno-risueno et al., 2016, Xuan et al., 2016]. In addition, studies have demonstrated an important role for synthesis of the auxin precursor indole-3-butyric acid (IBA) in the root cap (RC) for the amplitude of priming oscillations [Strader and Bartel, 2011, Xuan et al., 2015], and showed a reduced production of LRs for mutations in auxin transporting proteins such as PIN2 [Xuan et al., 2016], LAX3 [Swarup et al., 2008, Lewis et al., 2011] and AUX1 [Lewis et al., 2011, De Smet et al., 2007, Xuan et al., 2015]. Furthermore, auxin perception in the vasculature was shown to be critical for LR formation [De Smet et al., 2007]. Together, these studies indicate the importance of auxin production, transport and perception for LR priming. However, they do not explain the repetitive, oscillatory nature of the priming process. A recent study reported a strong spatio-temporal coincidence of repetitive apoptosis of the upper lateral root cap cells and priming events. Based on this correlation the authors proposed lateral root cap (LRC) apoptosis as the source of the oscillatory priming signal [Xuan et al., 2015, Xuan et al., 2016]. Nonetheless, *smb* mutants defective in LRC apoptosis still have LR formation [Xuan et al., 2016], indicating that this can not be the sole or true driver of priming. Still, these results may hint at a role for growth dynamics in the lateral root priming mechanism.

Based on the above findings, we decided to investigate how auxin transport and root growth dynamics together may impact LR priming dynamics, using a computational modeling approach. To this end we developed a new multi-scale root growth model in which we combine realistic root tip architecture and auxin transport, cell growth, division and expansion dynamics, and developmental zonation. Excitingly, our model shows that by these properties, repetitive elevation of auxin levels in the vasculature and pericycle automatically emerge. Specifically, the model shows that the architecture of the auxin reflux loop results in an auxin loading zone at the proximal end of the root tip meristem, with auxin preferentially being loaded in the narrow cells of the vasculature and pericycle. Root growth dynamics naturally produces regular alternations in the size of cells arriving at this auxin loading zone, which subsequently become amplified during cell expansion. This leads to substantial differences in the potential of cells to load auxin. The combination of these two properties results in regular fluctuations in the auxin levels of vascular and pericycle cells at the end of the meristem.

Our model predicts that the rate at which LR competent sites are formed predominantly depends on cell cycle duration, while the spatial patterning of LRs along the MR is controlled by cell cycle duration together with meristem size.

## 2 Material and Methods

### 2.1 General description of the model

We developed a novel multi-scale root growth model, combining from our earlier root models realistic root tip architecture [Van Den Berg et al., 2016] with root growth dynamics [Mahonen et al., 2014]. Like our earlier models, the model incorporates cell type specific (Fig 1A) and zonation dependent (Fig 1B) gene expression and polarity patterns of AUX/LAX auxin importers (Fig 1C) and PIN exporters (Fig 1D), developmental zone specific cellular growth, division, expansion and differentiation dynamics, cell level control of gene expression, and sub cellular, grid level, simulation of auxin dynamics. With respect to gene expression, the model incorporates the auxin-dependent gene expression of AUX/LAX.

**Figure 1:**
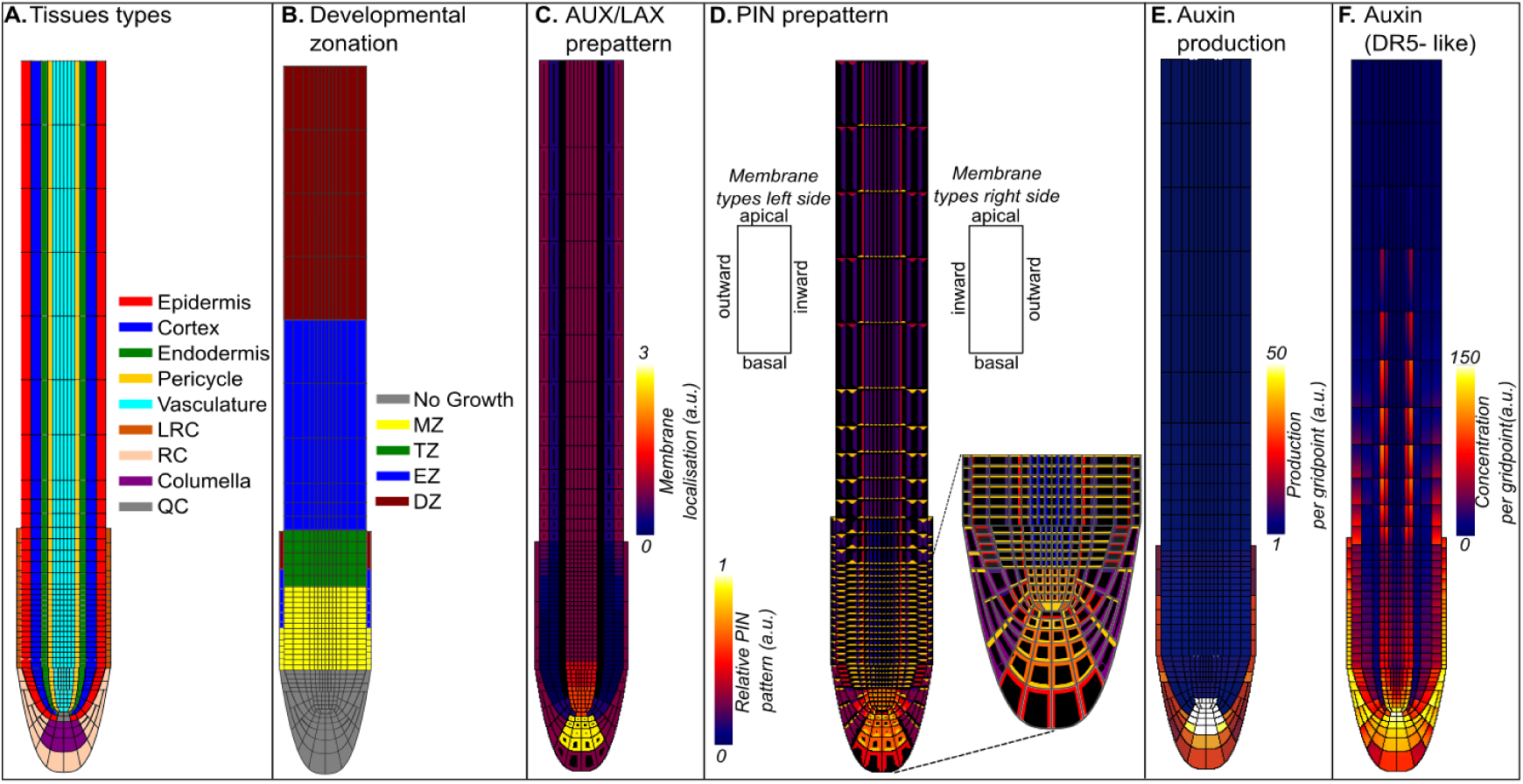
Model layout. **A**. Tissue types; columella (purple), RC (pink), LRC (orange), epidermis (red), cortex (blue), endodermis (green), pericycle (yellow), vasculature (cyan) and QC (gray). **B**. Developmental zonation; The root consisted of 4 zones: Meristem zone (MZ, yellow), transition zone (TZ, green), elongation zone (EZ, blue) and differentiation zone (DZ, red), the curved part of the root (grey) had no growth dynamics. **C**. Pre-patterned membrane localisation of AUX/LAX transporters, auxin feedback will shape the final membrane pattern. **D**. Predefined PIN polarisation pattern, with apolar PIN localization in columella and basal MZ, mainly apical PIN localization in epidermis and RC, vasculature and pericycle with predominant basal PIN localization, cortical PINs flipped from mainly basal to mainly apical upon EZ entrance. Inset is a magnification of MZ PIN polarisation pattern, all cells had similar PIN expression level. **E**. Auxin production level per cell, auxin production is limited to cytosolic gridpoints and cellular production is relative to cell size. **F**. Spatial distribution of auxin in equilibrium, DR5-like auxin reporter is used with a saturated hill-function

### 2.2 Tissue lay-out

In the current study we aimed to investigate the interplay between auxin transport and root growth dynamics. For this aim we need to realistically describe root tip auxin transport patterns, necessitating the incorporation of a wedge-shaped root tip layout encased in a lateral root cap (LRC). Our earlier research demonstrated the importance of such a realistic layout, as compared to a simplified rectangular root topology, for root tip auxin patterning [Van Den Berg et al., 2016]. At the same time, our research goal requires the incorporation of root growth dynamics. However, since the development of a full mechanical model of root growth dynamics is outside the scope of the present paper, the aim was to use the previously applied simplistic method of simulating root growth dynamics in which cells grow by adding a row of grid points and shifting upward all more shootward cells. While this root growth algorithm can be easily applied in a square root topology in which all cells are stacked in straight columns, this approach is less easily extended to the curved regions of the root tip. Therefore, as a compromise, we limited the size of the curved part of our root topology and ignored cell growth and divisions there, simulating growth dynamics only in the straight part of the root architecture. We reasoned that this is a justified approximation since it only ignores growth dynamics of the columella and lowermost parts of the RC, which growth dynamics are outside the scope of the current study, as well as the growth dynamics of the very slowly dividing quiescent center (QC), stem cell niche (SCN) and directly abutting cells.

The root layout was simulated on a grid of 224×1646 *μm*^2^ with a spatial resolution of 2 *μm*. Width of individual cell types was based on experimental data and earlier modeling studies [Laskowski et al., 2008, Van Den Berg et al., 2016]. A total of 8 different cell types were incorporated in the model. Moving from outermost to innermost these are: RC (pink), LRC (orange, 8 *μm*), epidermal (red, 18*μm*), cortical (blue, 20 *μm*), endodermal (green, 12 *μm*), pericycle (yellow, 8 *μm*) and 3 vasculature files (cyan, 6 *μm*). Finally, the vasculature converges on the QC (gray) and below the QC are the columella cells (purple) (Fig. 1A). The root was subdivided into 4 distinct developmental zones, moving from the root tip shootward these are: meristematic zone (MZ), with cytoplasmic growth and cell division; transition zone (TZ), with cytoplasmic growth but without further cell division; elongation zone (EZ), with vacuolar expansion; and differentiation zone (DZ), in which cells undergo terminal differentiation without growing further (Fig. 1B). In the model used in this study, to simplify matters, the position of zonation boundaries were defined in terms of distance from the QC rather than made dependent on auxin [Grieneisen et al., 2007] or PLT [Mahonen et al., 2014] gradients. Boundary positions were set such that the meristem proper has a size of 20 cells with an initial height of 8*μm*, the TZ has a size of 15 cells with an average height of 12 *μm*, the EZ was set at a size of 10 cells of 60 *μ*m average, and the DZ contained 5 cells of 100 *μm.* PIN expression and polarity patterns as well as AUX1/LAX patterns where incorporated based on tissue type and developmental zone, in agreement with experimental data [Bennett et al., 1996, Swarup et al., 2001, Swarup et al., 2005, Péret et al., 2012] and similar to earlier modeling studies [Grieneisen et al., 2007, Laskowski et al., 2008, Mahonen et al., 2014] (Fig. 1C and 1D). The pattern of the auxin transporters results in reverse fountain auxin flux pattern with maximum levels in the QC [Grieneisen et al., 2007, Mahonen et al., 2014, Van Den Berg et al., 2016](Fig 1F).

### 2.3 Auxin dynamics

Auxin metabolism, passive and active transport were implemented on a subcellular, grid point level in a similar manner as in earlier studies [Grieneisen et al., 2007, Mahonen et al., 2014, Van Den Berg et al., 2016]. Specifically, auxin transport was modeled as a combination of passive and active import into the cells, while auxin export was modeled as being purely active. Diffusion was modeled with distinct rates for cytoplasmic and apoplastic diffusion [Grieneisen et al., 2007, Mahonen et al., 2014, Van Den Berg et al., 2016] (Table 1). For a cytoplasmic grid point i,j (*A*_*i*, *j*_) surrounded by *n* wall (*A_wall_*) and m cytoplasmic (*A_cell_*) grid points the equation is as follows:

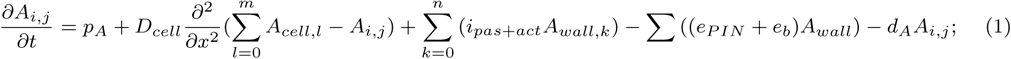

**Table 1:**
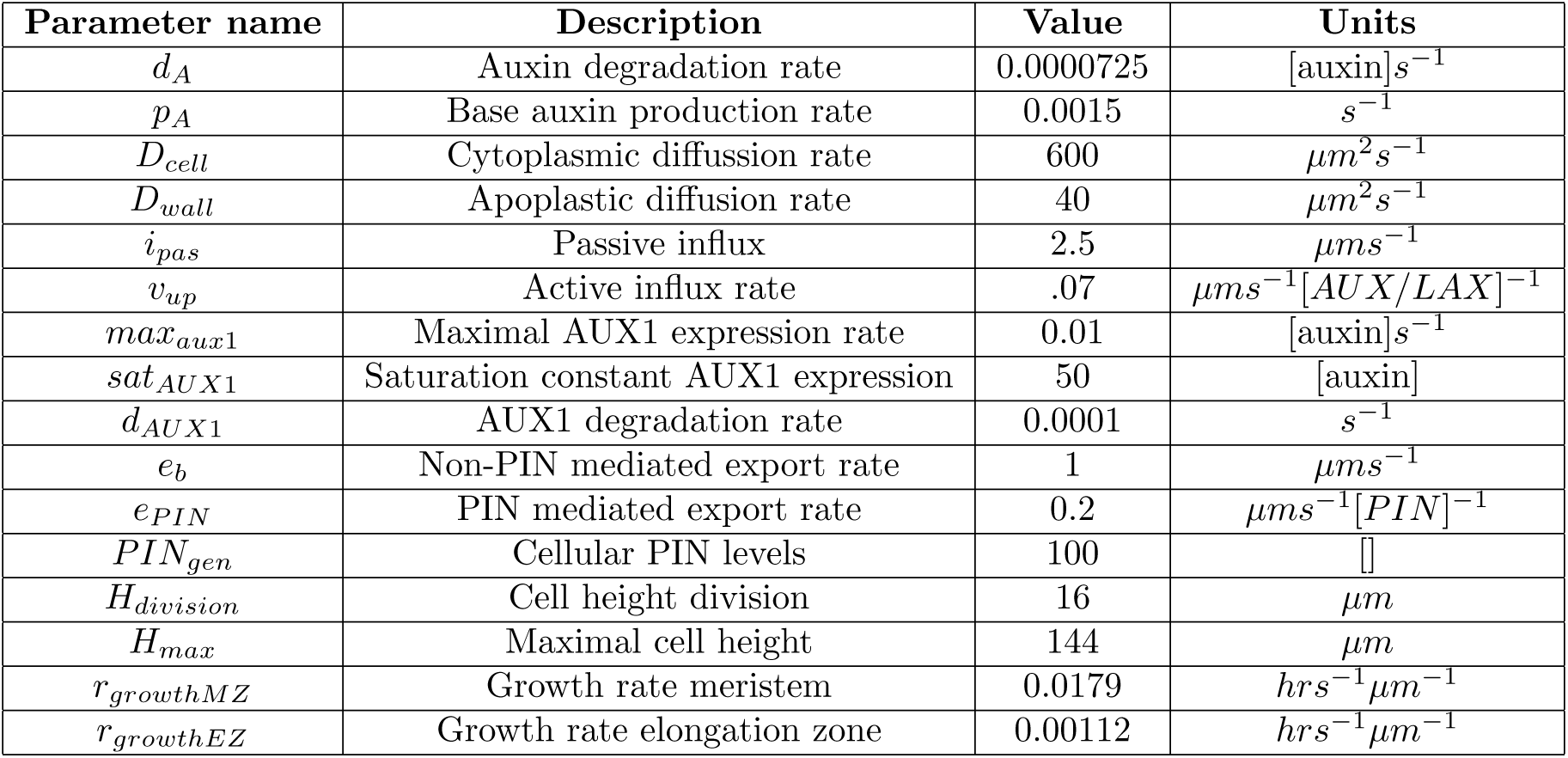
Model parameters used for default simulations

Here, *P_A_* is the auxin production rate, *d_A_* is the auxin degradation rate, and *D_cell_* is the diffusion rate of auxin inside a cell. *i*_*pas*+*act*_ is the combined passive, diffusional and active, AUX/LAX mediated influx of auxin from walls to cytoplasm, *e_PIN_* represents active, PIN mediated export of auxin from cytoplasm to walls, and active transport by other not explicitly modeled exporters such as ABCBs is captured in *e_b_.* For an apoplastic grid point i,j (*A*_*i*, *j*_) surrounded by *n* wall (*A_wall_*) and *m* cytoplasmic (*A_cell_*) grid points the equation is as follows:

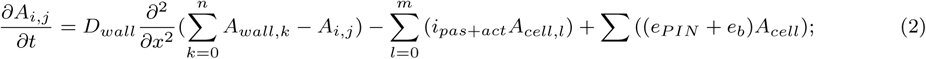

With *D_wall_* representing the auxin diffusion rate in the apoplast.

#### 2.3.1 Root tip auxin production

While historically, root auxin levels were assumed to almost solely depend on shoot delivered auxin, more recent data show the importance of root localised regions of high auxin production, particularly once roots have passed a particular developmental age [Bhalerao et al., 2002]. We incorporated elevated auxin production occurring in cells surrounding the QC as well as in the columella and LRC cells (Fig. 1E), assigning these cells with higher values of *pA.*

Finally, to ensure that despite grid based modeling of auxin production, the overall auxin production of an individual cell is independent of cell size we normalized *P_A_* as *P_A_*=p^∗^ *Height_cell_*/*Height_MZcell_*. Where *Height_cell_* is the height of the cell and *Height_MZcell_* is the initial height of a meristematic cell.

#### 2.3.2 AUX/LAX pattern

For simplicity active auxin import was described using a single AUX/LAX prepattern that represents the sum of all experimentally reported expression domains of AUX/LAX genes [Bennett et al., 1996, Swarup et al., 2001, Swarup et al., 2005, Péret et al., 2012] (Fig. 1C). Active AUX/LAX mediated influx is described as: *i_AUX/LAX_* = *v_up_* ^∗^ *AUX/LAX_pat_* ^∗^ *AUX/LAX_gen_*, where *v_up_* is the auxin uptake rate of AUX/LAX, *AUX/LAX_pat_* is the pre-patterned maximum membrane pattern of the auxin importers (Fig 1C) and *AUX/LAX_gen_* is the cell level gene expression of AUX/LAX. AUX/LAX expression is auxin dependent [Laskowski et al., 2006, Laskowski et al., 2008], and we recently showed that this auxin dependence plays an important role in root tropisms [Van Den Berg et al., 2016]. Here we assume that auxin has a saturated positive effect on AUX/LAX cellular expression levels in the following way:

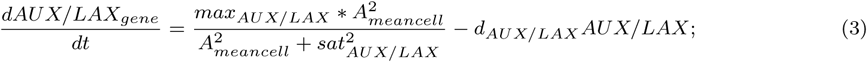

Here, *max_aux_/_lax_* is the maximal gene expression rate of AUX/LAX, *sat_AUX/LAX_* is the auxin level at which the rate of AUX/LAX expression is half maximal, AUX/LAX proteins are degraded with rate *d_AUX/LAX_*, and *A_meancell_* is the average cellular auxin level

#### 2.3.3 PIN expression and localisation

Similar to our earlier studies, we model active auxin export from cells as consisting of a major PIN protein mediated component (*e_pin_*) and a minor additional component (*e_b_*) that can be thought of as ABCB/PGP mediated auxin export. For simplicity (*e_b_*) is assumed equal for all cells and have an apolar cellular pattern. With regards to PIN mediated transport, tissue type and zonation dependent PIN pre-patterns are incorporated based on experimental data and similar to those used in earlier models (Fig 1D) [Grieneisen et al., 2007, Laskowski et al., 2008, Mahonen et al., 2014, Van Den Berg et al., 2016]. Relative to earlier models changes were made in the PIN1 polarity pattern in the MZ, based on recent experimental data demonstrating a relatively apolar distribution of PIN1 in the lowermost regions of the root [Omelyanchuk et al., 2016](Fig. 1D). This change resulted in a broader, more robust auxin maximum, more consistent with experimentally observed auxin patterns (Compare Fig. 1F and S1C).

### 2.4 Growth dynamics

Earlier data on Arabidopsis root growth dynamics [Beemster and Baskin, 1998] suggested that cell cycle durations in the root apical meristem (RAM) are in the order of 20 hours. These cell cycle durations were based on measured cumulative cell flux dynamics at the end of the meristem with the assumption that all, approximately 30-35, rows of cells within the meristem divide at a similar rate. In our earlier model, cellular growth dymamics were based on these estimated rates [Mahonen et al., 2014]. However, more recent data suggest that cell divisions occur in only a limited, rootward region of the meristem containing 15-20 cell rows. Cells in the remaining more shootward part of the meristem grow slowly, while not or hardly dividing, until switching to rapid vacuolar expansion driven growth in the elongation zone [Dello Ioio et al., 2008, Novák et al., 2016]. Division rates measured within the lowermost, actively dividing part of the meristem were found up to 3 hours per cell cycle [Campilho et al., 2006, von Wangenheim et al., 2017]. To account for these recent insights, we incorporated in the current model maximum cell division rates of 7hrs. In addition, we also explicitly incorporated a proper meristem zone (MZ) in which cells actively divide and a shootward MZ part, which we will refer to as a transition zone (TZ), and in which we ignore rare cell divisions and only simulate slow cytoplasmic cell growth (Fig 1B).

Individual cells start in the MZ where they grow and divide with rate *r_growthMZ_*/*μm* and divide when they have doubled their size. Noteworthy, in our model several rows of cells that are part of the meristem do not actually divide due to their residence in the curved part of the root tip and our simplified growth algorithm not being applicable there (see Tissue Layout section). To check whether this influences priming dynamics we performed simulations where the curved region of the root tip was decreased further in height, containing only the QC and SCN, and with all other MZ cells now being able to grow and divide. We found that this did not qualitatively affect priming dynamics (Fig S2C). When leaving the MZ, cells enter the TZ where they still grow with *r_growthMZ/μm_* but no longer divide. Upon entering the EZ, cells start to expand with rate *r_growthEZ_*/*μm* until a maximum cell height is reached and cells enter the DZ. MZ and EZ growth rates are per /*μm*, resulting in higher per cell growth rates for larger cells and constant elemental growth rates, consistent with experimental observations.

Given the discrete, grid based nature of our model, cellular growth is executed in discrete steps during which a single row of grid points is added to the height of a cell. The time interval at which these discrete growth event occurs follows from the cellular growth rate in the following manner *if*(*time*) ≤ *time_prevgrowthstep_* +(1./(*r_growthMZ/EZ_* ^∗^ *cellheight*)) add row of gridpoints. Concentrations of auxins and proteins are corrected for these instantaneous cellular volume increases in case of cytoplasmic growth, but not in case of vacuolar driven cell expansion where cytoplasmic volume is assumed to stay constant. Upon division, cells are divided into two equally sized daughter cells that inherit transporter patterns and concentrations of cellular components of their mother cell. All tissues grow in the described manner, except for LRC tissue which has shorter developmental zones, lacks a TZ, and where cellular apoptosis occurs when cells reach a fixed position from the root tip, corresponding with the start of EZ of other tissue types (Fig. 1B).

To ensure that the simulated tissue stays within the range of the simulation domain, the most apical cells are removed if their apical cell wall is within 2 grid points of the simulation domain boundary.

### 2.5 Model variations

#### 2.5.1 Auxin dynamics

Unless explicitly stated differently, simulations were performed with the above described settings for auxin dynamics, with parameter values as described in Table 1. However, to investigate the impact of auxin transport through the root tip reflux loop and the effect of root tip auxin metabolism, several simulations were done with altered expression and/or localization of auxin importers and exporters or altered auxin production. Alterations in transporter levels or auxin production rates were applied by simply multiplying default levels with a parameter *α*, using *α* > 1 in case of increase and *α* < 1 in case of decrease of transporter or production levels. Alterations were often applied in a tissue and zone specific manner, applying *α* ≠ 1 only in specific regions of the root tip.

#### 2.5.2 Normalizing auxin fluxes and pH effects on background influx

To investigate the influence of expansion driven increases in cell size on cellular auxin uptake and release dynamics we performed simulations in which we normalized auxin uptake and release with cell size.

Auxin import, be it passive or active, increases with cell size. Passive transport occurs across the membrane, and membrane surface area increases with cell size. Additionally, since active AUX/LAX transporters have an apolar cellular pattern, and in our default simulations neither gene expression levels nor prepatterns are rescaled with cell size, active transport is also implicitly assumed to increase with cell surface area and hence size. To investigate the relevance of these increases in passive and active auxin import with cell size, we normalized these auxin fluxes by multiplying them with a factor 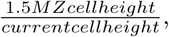 where 1.5*MZcellheight*(= 16*μm*) is the average cell height in the TZ just before expansion starts, which is 1.5 times the size of just divided MZ cells, and *currentcellheight* is the actual cell height. We thus effectively dilute the auxin fluxes as cell size increases, which in case of active uptake may be a more reasonable assumption.

In contrast, PIN transporters have a predominant polar localisation on the non-growing apical (PIN2) or basal (PIN1) membranes. As a consequence, as cell size increases auxin export decreases relatively. To check the consequences of this relative decrease in auxin export specifically for vasculature cells, we multiplied basal PIN1 levels with a factor 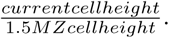 Note that this is the inverse of the factor used above, as now we aim to increase rather than decrease auxin transport with cell size to compensate for its decrease with cell size. All normalisations were only applied for cells with a height exceeding 1.5*MZcellheight*, and were not applied to lateral root cap and columella cells.

Additionally, to study the influence of passive auxin uptake, rather than normalizing it for cell size, we implemented a modest pH dependent increase of background influx in the EZ [Rutschow et al., 2014, Street et al., 2016, Barbez et al., 2017]. For this we implemented a tissue pH gradient, with constant high pH levels in the MZ (*pH_MZ_* = 5.8) and a steep drop of pH in the EZ (minimum pH in EZ of 4.8, reached at a distance of 150 *μm* from the start of the EZ), and a recovery of the same high pH levels in the DZ (reached directly at the start of the DZ). To couple pH levels to passive influx levels we applied the following formula:

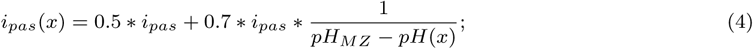

Were the *i_pas_* is the default background auxin influx rate used thusfar (Table 1) and the baseline passive influx was set to 50% of *i_pas_* to compensate for the extra influx occurring in the EZ. The longitudinal position of a grid point along the root was represented by *x.* The high MZ and DZ pH levels result in only baseline passive influx, lower pH levels increase passive influx.

#### 2.5.3 Growth dynamics

Unless explicitly stated differently, growth dynamics were applied as described above. Under these default growth dynamics, all cells deterministically grow and divide at the same rate independent of tissue type or position in the MZ (apart from the dependence of growth rate on cell size). Similarly, all cells transit from the MZ to EZ at the same longitudinal position, independent of tissue type. As a consequence, cell growth and division dynamics are perfectly synchronized within and across cell files.

In reality, cellular growth and division dynamics are not entirely deterministic resulting in variation in growth rates and division sizes. To mimic stochastic variations in growth rate and cell cycle duration between cells, we performed additional simulations. We assigned growth rates that were drawn from a random distribution with a mean equal to the default growth rate and a maximum of either 10% or 75% deviation from this mean growth rate in both directions for each new transit amplifying cell arising from the division of the lowermost, stem cell like cells. To maintain a symmetric auxin profile and hence tissue architecture, identical growth rates were applied to pairs of cells occupying the same position relative to the root mid line. Also, we performed simulations in which cell cycle duration was constant and homogeneous within a cell file, but different between cell files of different cell types. Cell cycle durations ranging from 5.2 to 8.5 hours were chosen for the different tissue types. Again, cell cyle durations of cells occuyping mirror image positions relative to the midline were taken the same to maintain symmetric growth.

In addition, in real plants cellular growth and division rates depend on auxin levels, which, given the auxin gradient extending from the QC, results in a gradually declining growth rate profile. To simulate this, further additional simulations were performed in which such a growth rate distance profile, intended to emulate the auxin dependence of cell cycle duration, was implemented. To achieve this we use the function *r_growthMZ_/dist*, with *dist* being a increasing function of distance to TZ. Finally, in *Arabidopsis* it can be seen that while vasculature cells start expanding relatively close to the root tip, pericycle cells transit to expansion furthest from the root tip. Thus, in a final set of simulations we implemented a tissue type dependent location of the MZ-TZ boundary.

#### 2.5.4 Local growth simulations

Apart from the curved part of the root model, under default conditions all cells exhibit zonation dependent growth dynamics. To gain a better understanding of the role of growth induced cell size increases on auxin levels, artificial growth simulations were performed in which only a limited set of cells were allowed to grow. For this first a default growth simulation was run until a steady state was reached in which auxin concentration and gene expression levels no longer significantly changed. This steady state situation was subsequently used as an initial condition for the artificial, localised growth simulation. A first set of simulations was performed in which only 1 row of cells was allowed to grow, varying the location of this row of cells along the length of the root. The goal of these simulations was to investigate the role of cell size increase and cell location on auxin dynamics (used in Fig 6C, D and E).

We performed an additional set of simulations, the goal of which was to determine the independent influence of residence time, cell size and size of the below neighboring cell on the amount of auxin loading occurring at the start of the EZ. As before, these artificial growth simulations were started from steady state conditions obtained under normal growth dynamics. For these simulations, controlled growth was applied as follows: Only the upper 10 cells of the MZ were allowed to grow. In addition, linear growth was applied, meaning that cellular growth rates do not increase with cell size as is normally the case. These specific growth dynamics were maintained irrespective of whether cells are still in the MZ or enter the EZ. The non-standard linear growth dynamics were chosen for more easy control of growth rate and hence overall cumulative displacement generated by this growth domain as opposed to the standard exponential growth. Cells that are in the EZ from the start of the simulation onwards elongated with standard, exponential dynamics.

First, to investigate the effect of loading time we varied the growth rate in the MZ, essentially
varying displacement speed. However, without additional measures, changing the time spent in the early EZ will also affect the growth time and hence the size the cell has when residing in this zone. To be able to investigate the impact of residence time independent of cell size, elongation rates need to be adjusted such that a constant final cell size is reached at a fixed distance from the root tip. To achieve this, cells in the EZ were tracked and their actual height, *height_act_* is compared with the target height, *height_tar_* that would normally be achieved under default growth rates. The ratio between these two heights is next used to determine a modified root growth rate:

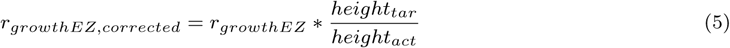

This corrected growth rate is subsequently applied, resulting in *height_act_* converging to *height_tar_*. In this way growth rate variations in the below MZ are compensated by changes in the growth rate in EZ. The target cell height is determined as follows: *height_tar_* = *height*_start_+(*r_growthEZ_*)^*t_growth_*^, where *height_start_* is the cell size at the start of the simulation and *t_growth_* is the time a cell would need to reach the current position under default displacement velocity. In addition to varying loading time, we also investigated the effect of the size of the below neighboring cell. For this we simply varied whether or not the below MZ cells divide. Finally, we tested the effect of the size of the cell itself by varying the expansion rate, *r_growthEZ_*, in the EZ while keeping the final to be obtained size constant(Fig 8).

#### 2.5.5 LRC mutants

In simulations of wild type plants, cells of the LRC are simulated to undergo apoptosis and shed off when the basal membrane of the top LRC cell passes the border of TZ and EZ of the epidermis. To investigate the influence of LRC apoptosis on auxin oscillations we mimicked a *fez* and *smb* mutant, that have delayed or premature LRC shedding respectively [Willemsen et al., 2008](Fig. 4B and C). To mimic a *fez* mutant we assumed premature LRC shedding to occur at a position just above the start of the growing part of the root, essentially allowing for only a minimum of LRC cells to occur(Fig. 4B). For the *smb* mutant we assumed that LRC shedding only occurs when reaching the end of the simulation field, corresponding to a position beyond the start of the differentiation zone (Fig. 4C).

### 2.6 Numerical integration and run-time performance

Auxin dynamics were solved with a temporal integration step of Δ*t*=0.2s and a spatial integration step of Δ*x* = 2*μm*. Importantly, auxin transport occurs at relatively high rates. As a consequence, standard Euler forward explicit integration schemes would require very small temporal integration steps (Δ*t*=0.0001) to solve the above equations stably. Given that we aim to simulate plant growth dynamics over a time course of several days, this would result in excessively long simulation run times. Therefore, similar to earlier modeling studies [Grieneisen et al., 2007, Mahonen et al., 2014] we used an alternating direction semi-implicit integration scheme, allowing us to use integration steps of 0.2s. Simulations were run on a desktop with E5-2687Wv4 processor. The code of the model was written in C++ and typical run-time was 24 hours, for a simulation representing 6 days of plant growth.

### 2.7 Analysis Methods

To display the spatial and temporal dynamics of the observed auxin oscillations in one single plot, we generated space time plots. These kymographs (space time plots) were created by taking snapshots of the auxin patterns inside the cells as well as the locations of cell walls in a one grid point wide longitudinal cross section in the tissue of interest. For this we typically used the average cellular auxin level of the outer vasculature tissue file. In terms of spatial extent, space time plots started at the first dividing cell ( 250 *μm* from root tip) and ended when cells are fully elongated ( 1050*μm* from the root tip), essentially capturing all growing tissue in one longitudinal shot. Snapshots were stored every 100 time steps (=20s) and aligned according to their temporal sequence (Fig 2, FigS3).

**Figure 2:**
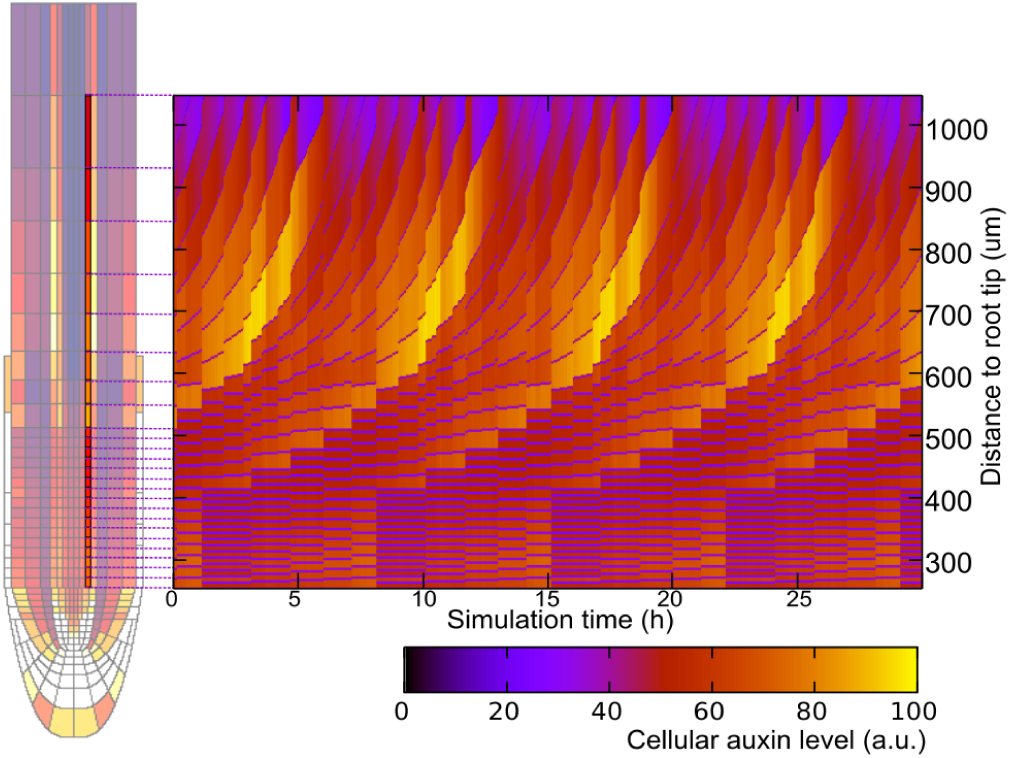
Explanation of Kymograph. Kymographs (space time plots) were created by taking every 100 time steps (20 sec.) an one grid point wide longitudinal snapshot of the growing region in the root and aligning these sequentially. Snapshots were taken from outer vasculature cell file and cellular auxin levels were on the indicated scale, unless explicitly stated differently. From the kymograph it becomes clear that temporally regular peaks of auxin arise near the end of the MZ ( 600 μm from the root tip).

### 2.8 Robustness of the model

To test whether the observed behavior found here might be an artifact of modeling choices we performed an extensive robustness analysis. We tested the influence of auxin production rates and locations, details of auxin importer and exporter spatial patterns and gene expression dynamics, size of the overall simulation field, spatial resolution and root architecture (see Supplemental Results). We found that none of these variations in model settings abolishes the observed auxin oscillations.

## 3 Results

### 3.1 Auxin oscillations as an emergent property of the new model

Here we aimed to investigate the contribution of root tip architecture, auxin reflux loop topology and root growth dynamics on lateral root priming. To this end we developed a novel multi-scale root model incorporating all these properties (see Methods). Notably, previous models have at best either incorporated a realistic root topology [El-Showk et al., 2015, Van Den Berg et al., 2016, Di Mambro et al., 2017] or root growth dynamics [Grieneisen et al., 2007, Mahonen et al., 2014], but none has thusfar combined these two. Surprisingly, with these properties incorporated, regular temporal variations in vascular and pericycle auxin levels automatically emerge in our new model (Fig 2, Fig S3, movie S1). To confirm that the observed oscillations represent a genuine auxin oscillation mechanism, rather than an artifact of model assumptions or implementation choices, we performed a range of alternative simulations. These simulations included variations in the precise root architecture, length of the simulation field, spatial resolution of our simulations, as well as auxin production rates and locations and auxin transporter patterns that all displayed similar oscillatory dynamics (Supplemental Results 1, Fig S1 and S2).

### 3.2 Auxin reflux loop determines auxin loading domain

To unravel the mechanism underlying the observed auxin oscillations, we first investigated the impact of auxin transport. PIN mediated auxin fluxes can roughly be divided into 3 domains. First there is the downward transport mediated by the basally positioned PIN1, PIN3 and PIN7 in the vasculature and endodermis, next there is inward transport mediated by laterally positioned PIN2 in the distal meristem and early EZ, and finally there is the shootward transport mediated by the apolarly localized PIN3, PIN4 and PIN7 pumping auxin out of the columella and the apically oriented PIN2 transporting auxin upward through the LRC and epidermis.

To investigate the importance of these distinct parts of the reflux loop we performed several simulations. First, we performed a simulation in which we removed PIN2 from the epidermis, cortex and LRC as well as PIN3, PIN4 and PIN7 from the columella, leaving only PIN1 in its normal expression domain as well as PIN3 and PIN7 in their vascular domain. This resulted in a nearly complete abolishment of auxin oscillations (Fig. 3A). Next, we performed a simulation in which laterally positioned PIN2 were only allowed to occur inside the MZ, but not in the TZ or higher shootward. This again resulted in a reduction of auxin oscillation amplitude, albeit considerably less severe (Fig. 3B). Finally, we simulated auxin dynamics in the presence of normal PIN1, PIN3, PIN4 and PIN7 levels and patterns, but with strongly reduced PIN2 levels (reduction of 67%) while maintaining PIN2’s normal polarity pattern. Again, auxin oscillation amplitude is reduced, but to a lesser extent as when PIN2 is specifically reduced laterally (Fig. 3C).

**Figure 3:**
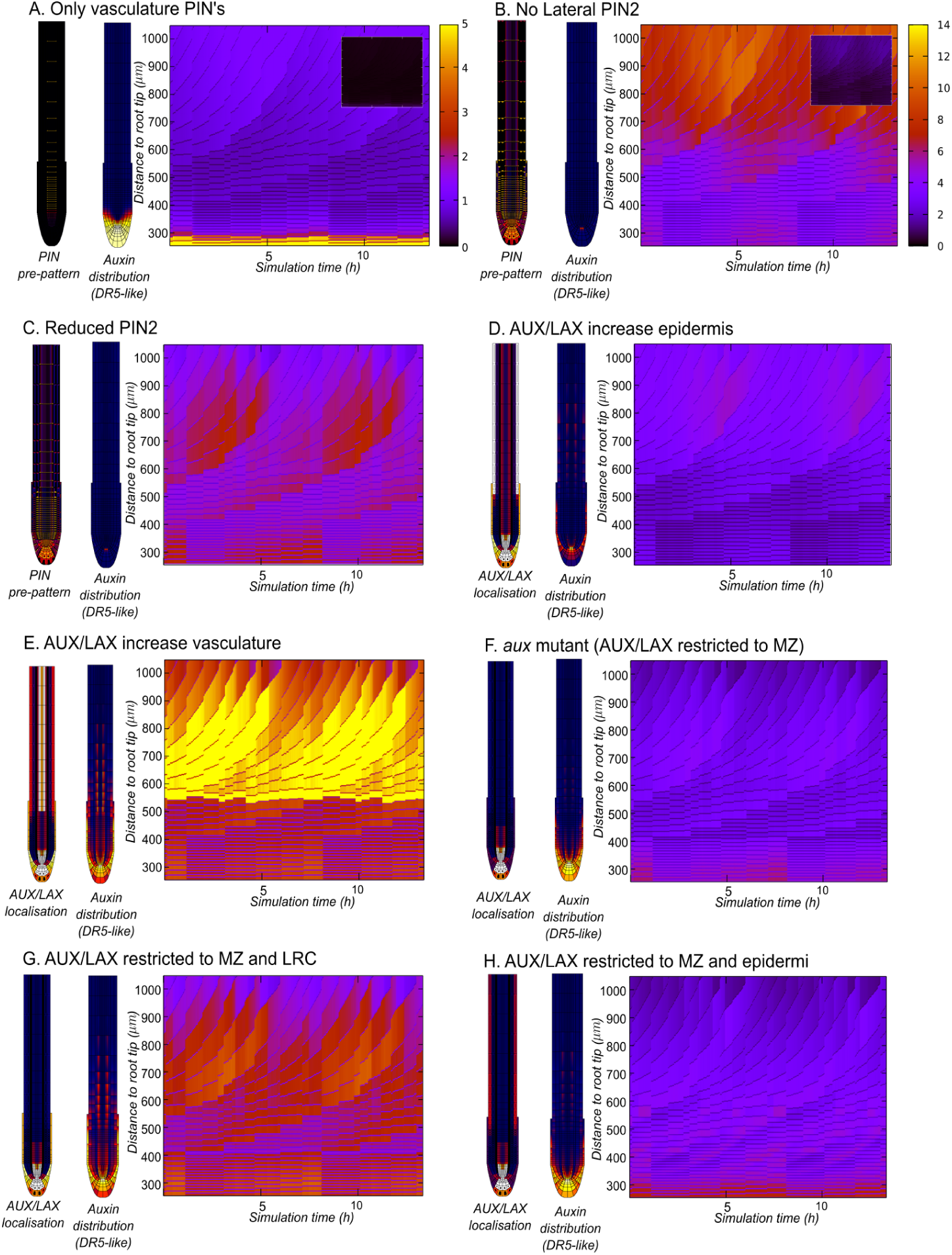
Auxin reflux loop properties create an auxin loading zone at the TZ and start EZ. **A**. Root with only PIN1 expression, auxin levels strongly decrease and oscillations vanish (Kymograph has adapted scale for visual reasons, inset shows kymograph with default scale). **B**. No lateral PIN2 in TZ and EZ, auxin levels and oscillations strongly decrease (Kymograph with adapted scale, inset shows default scale). **C**. PIN2 in TZ and EZ is reduced with 67% while maintaining polarity patterns. Auxin levels and oscillations decrease, however less than in case figure A or B. **D**. Increased (factor 5) AUX/LAX expression in epidermal TZ/EZ, auxin accumulates in epidermis and oscillation amplitude strongly decreases. **E**. Increased AUX/LAX (factor 5) expression in vasculature tissue, auxin accumulates in vasculature and oscillation amplitude increases. **F**. aux-like mutant with only AUX/LAX expression in MZ, auxin levels and oscillations strongly decreases. **G**.aux-like mutant with AUX/LAX expression in MZ and RC, auxin levels and oscillation are partly restore13compared to F. **H**. textitaux-like mutant with AUX/LAX expression in MZ and epidermal EZ/TZ, auxin levels and oscillations are strongly decreased comparable to F.

The differences in severity of reduction of oscillation amplitude gave insight in the importance of a functional reflux loop. Auxin needs to be transported outward from the QC region by PIN3/PIN4/PIN7 in the columella region, upward to the TZ and EZ by apical PIN2 in the LRC and epidermis of the MZ, and inward to the vasculature by lateral PIN2 in the RC and epidermis of the TZ and EZ. In absence of PIN2 and PIN4 and columellar PIN3 and PIN7 hardly any auxin arrives at the TZ and EZ, almost fully abolishing oscillations. In absence of lateral PIN2 in the TZ and EZ auxin does arrive at the TZ, yet little auxin is subsequently transported into the vasculature, strongly reducing oscillations. Simply reducing PIN2, while maintaining ratio’s between apical and lateral membranes maintains the essence of first upward transport towards the TZ and then inward transport into the vasculature and hence has less severe effects on oscillation amplitude.

To validate this hypothesis of a reflux-loop defined auxin loading zone in the priming region, we investigated the effect of increasing AUX1 mediated auxin import in different regions and cell types. Based on our hypothesis we predict that increasing AUX1 levels in the vasculature of the TZ and EZ will draw more auxin there, thus elevating oscillation amplitude, while increasing AUX1 levels in the epidermis and LRC will retain auxin there, thereby reducing oscillation amplitude. Simulation outcomes agree with these predictions, thereby supporting our hypothesis (Fig. 3D and E).

Still, previous experimental studies have demonstrated that *aux1* mutants have lower amplitude auxin oscillations that can be restored by LRC specific but not EZ epidermal *AUX1* expression [De Smet et al., 2007]. Thus, while our simulations suggest that increased AUX1 in the LRC negatively affects auxin oscillation amplitude, these experimental data indicate that LRC AUX1 is required for sufficient oscillation amplitude. To investigate this seeming paradox, we simulated both *aux1* mutants and mutant plants conditionally expressing AUX1 in either the LRC or EZ epidermis (Fig. 3F, G and H), reproducing the experimental results. Thus, apparently sufficient LRC1 AUX1 is essential to guide auxin into the LRC from which it subsequently moves into the priming zone, yet high LRC1 AUX1 prevents the auxin from leaving the LRC. Combined this causes a bell-shaped dependence of oscillation amplitude on LRC AUX1 levels. Together our results indicate that the reflux loop architecture generates an auxin loading zone in the vasculature of the TZ and EZ that is essential for generating auxin oscillations.

### 3.3 Auxin availability modulates oscillation amplitude

It has been previously shown that the production of the auxin precursor IBA in the LRC and its subsequent transformation into auxin positively influences the amplitude of priming oscillations [Strader and Bartel, 2011, Xuan et al., 2015].Therefore, as a next step we reduced the high auxin production in the LRC. Consistent with experimental results, oscillation amplitude was reduced (compare Fig 2 with Fig. 4A). Importantly, reducing local auxin production in the region around the QC and SCN, as well as reducing the influx of auxin from the shoot, produced similar reductions in oscillation amplitude (Fig. S4). This indicates that the total amount of available auxin in the reflux loop rather than the precise origin of the auxin is relevant for oscillation amplitude.

**Figure 4:**
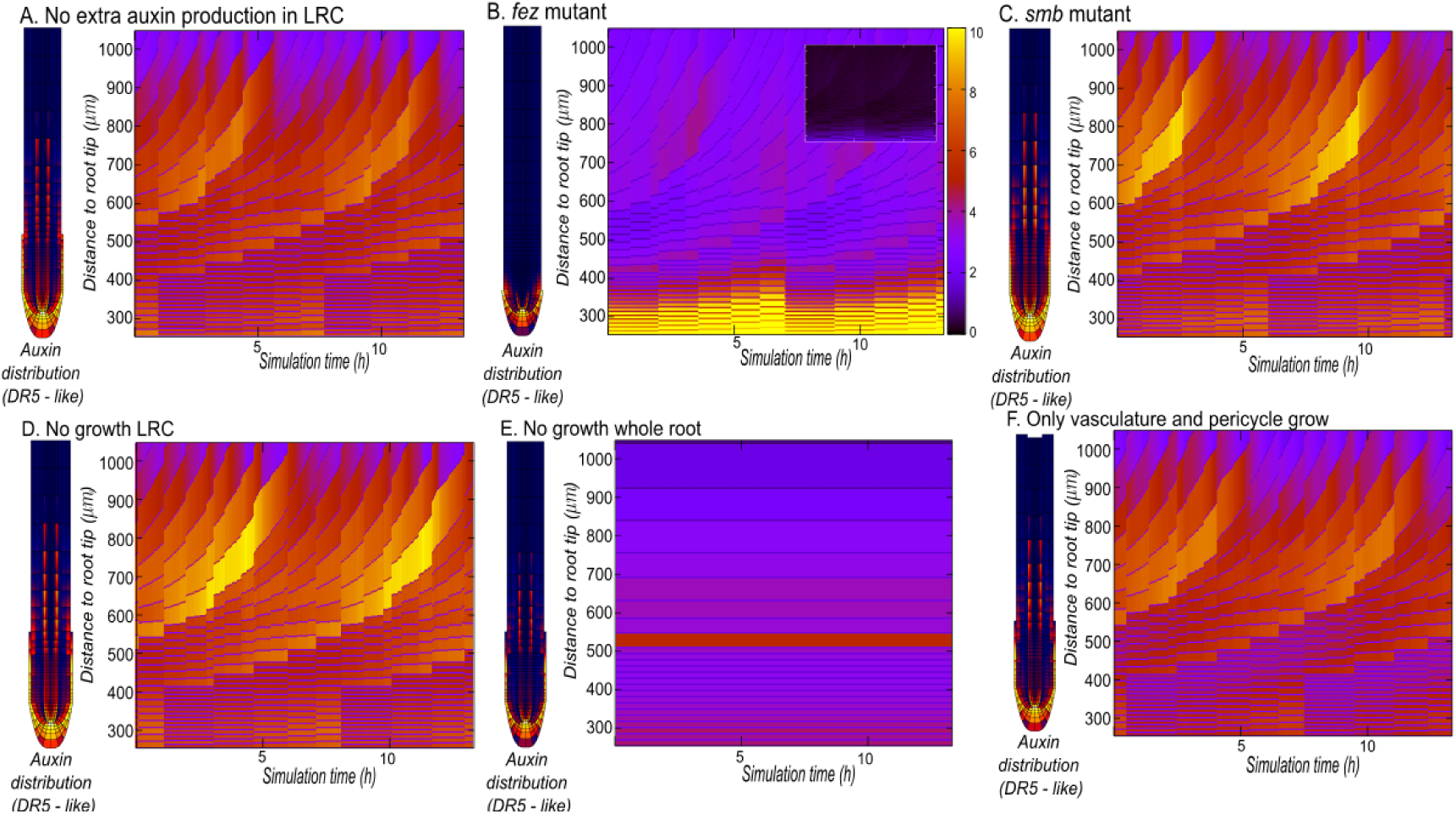
Vasculature growth and LRC auxin production and size are important for priming dynamics. **A**. Decreased auxin production in LRC, oscillation amplitude slightly decreases. **B**. fez mutant with premature LRC shedding, auxin accumulates in root tip and oscillation amplitude vanishes (Kymograph with adapted scale, inset shows default scale) **C**. smb mutant with defective LRC apoptosis, auxin levels and oscillation amplitude does not change. **D**. LRC does not grow or shed off, auxin level and amplitude hardly changes. **E**. Root without growth dynamics, auxin oscillations in vasculature are absent. **F**. Only vasculature and pericycle tissue files exhibit growth dynamics, oscillations are restored with comparable amplitude.

### 3.4 Non-LRC growth dynamics critical for auxin oscillations

Another recent experimental study suggested periodic LRC apoptosis as a mechanism for repetitive elevated auxin delivery that could drive oscillatory LR priming [Xuan et al., 2016]. To investigate the influence of LRC architecture as well as LRC growth dynamics, we simulated a *smb* mutant (extended LRC up into the DZ), a *fez* mutant (extremely short LRC), and a wild type (WT) plant root, in presence as well as absence of LRC growth dynamics (growth, division, expansion, apoptosis and shedding) (see Methods). We found that auxin oscillations were markedly reduced in amplitude in the *fez* mutant (Fig. 4B), but unaffected in the *smb* mutant (Fig. 4C). In addtion, growth of the LRC was not necessary for auxin oscillations (Fig. 4D). This indicates that for oscillations to occur, a minimum but not a maximum LRC length is needed. Furthermore, LRC apoptosis is not necessary for auxin oscillations, consistent with the observation that in the *smb* mutant only priming frequency and regularity are mildly affected [Xuan et al., 2016]. Combined with our previous findings from manipulating auxin transporters these results can be interpreted as follows: For oscillations to occur auxin needs to be supplied to the vasculature of the TZ. The LRC is a crucial part of the reflux loop as it strongly contributes to the early shootward and subsequent lateral inward transport, but can only function in delivering auxin to the loading zone if it extends up until this zone.

Next we investigated the role of growth dynamics in other tissue than the LRC. To do so we restricted growth dynamics to a subset of cell files. First, we simulated a root without any growth dynamics, resulting in the complete abolishment of the auxin oscillations(Fig. 4E). Since the priming signal occurs in the vasculature and pericycle tissue we proceeded by limiting growth dynamics to these tissues. We find that auxin oscillations are restored, with a similar frequency as observed in a root with growth dynamics in all cell files(Fig. 4F). The reduced amplitude indicates the importance of growth of neighbouring tissue for auxin availability but not for the appearance of the oscillations. Alternatively, when growth dynamics were limited to epidermis, cortex, endodermis or a combination of these tissues no auxin oscillations were observed(Fig. S5).

### 3.5 Tissue specific growth dynamics enable priming multiple pericycle cells

An apparent distinction between our simulations thus far and priming in planta is that in our model auxin maximum occur in a single pericycle and vasculature cell. In contrast, in planta, priming is thought to involve multiple pericycle cells, and subsequent competition amongst these cells prevents the formation of multiple nearby LRs from a single prebranch site [Dubrovsky et al., 2006]. Importantly, thusfar in our simulations cells in different cell files grow, divide, expand and differentiate synchronously. In real plants this is clearly not the case, cell divisions occur scattered among different cell files [Rahni and Birnbaum, 2018], and while vasculature cell files appear to expand and differentiate the closest to the root tip the abutting pericycle cells undergo expansion and differentiation furthest from the root tip, resulting in large vasculature cells next to small pericycle cells at the start of the TZ [Campbell and Turner, 2017].

First, we investigated the impact of asynchronous divisions between cell files, and found that this does not have significant effect on priming dynamics (see Methods) (Fig S7A). Next, we investigated the consequences of incorporating tissue specific positions of the TZ boundary, with vasculature cells transitioning closest and pericycle cells furthest from the QC. Incorporating these characteristics into our model results in an auxin elevation first occurring in a single large vasculature cell (Fig. 5A), as before, and subsequently being passed on to several pericycle cells (Fig. 5B). Additionally, it can be observed that the early onset of differentiation in the vasculature leads to an increased amplitude in oscillations (Fig. 5A). Thus, by incorporating these additional details, priming dynamics in our model even more closely mimic in planta observations. oscillation amplitude (Fig. S4). This indicates that the total amount of available auxin in the reflux loop rather than the precise origin of the auxin is relevant for oscillation amplitude.

**Figure 5:**
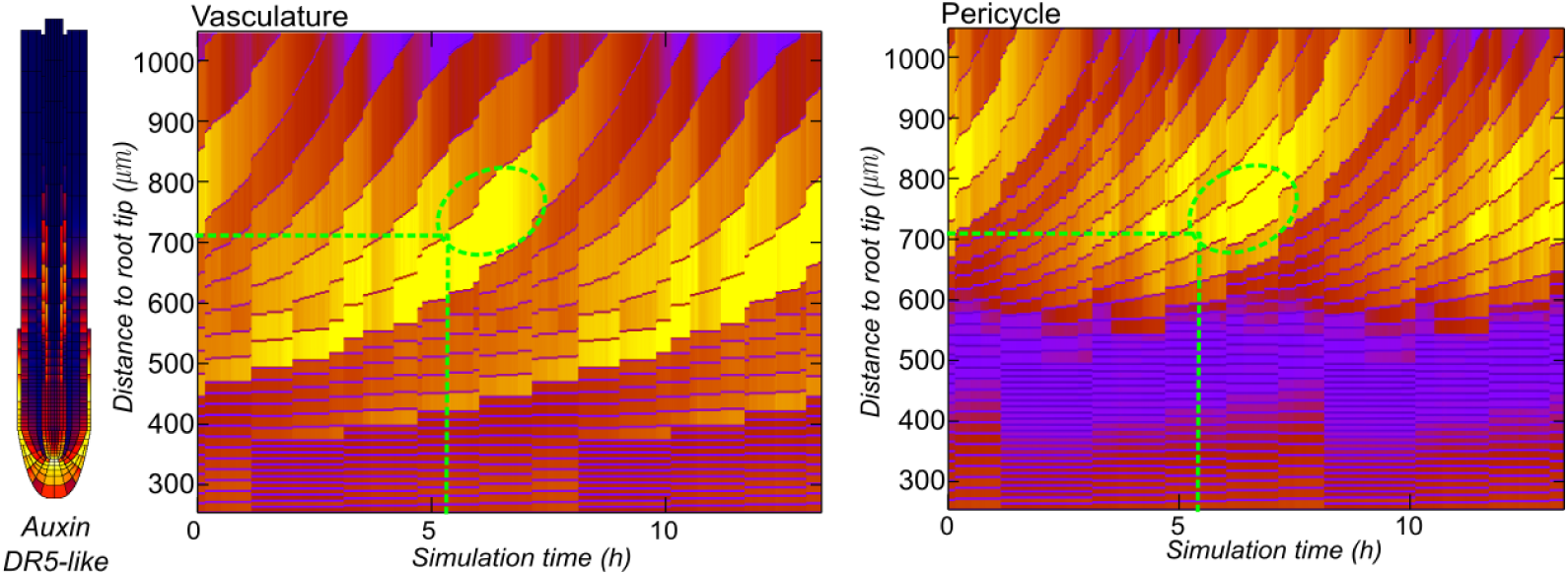
Priming of multiple pericycle cells. Early (more rootward) differentiation of the vasculature increases priming amplitude (Compare Fig. 2 with **A**.), additionally late (more shootward) differentiation of the pericycle gives a priming signal in multiple pericycle cells overlaying the single primed vasculature cell (**B**.). Green lines and circle are for comparing time and distance from root tip in the different tissues at the time of the auxin maximum.

### 3.6 EZ Vasculature cells have optimal auxin loading capacity

A close inspection of the kymographs suggests that the periodic changes in auxin levels we observe correlate with variations in the sizes of cells arriving in the TZ/EZ. To further validate this observation, we next plotted a kymograph using a color code for *cell size* rather than auxin levels and compared this to the default kymograph depicting auxin levels (Fig. 6A and 6B), confirming this idea. To determine whether there is a causal relation between cell size and auxin content we performed a simulation in an overall non growing root, applying only localized growth to a single row of cells while varying the longitudinal location along the root of these cells (see Methods section on Local growth simulations)(Fig. 6C).

**Figure 6:**
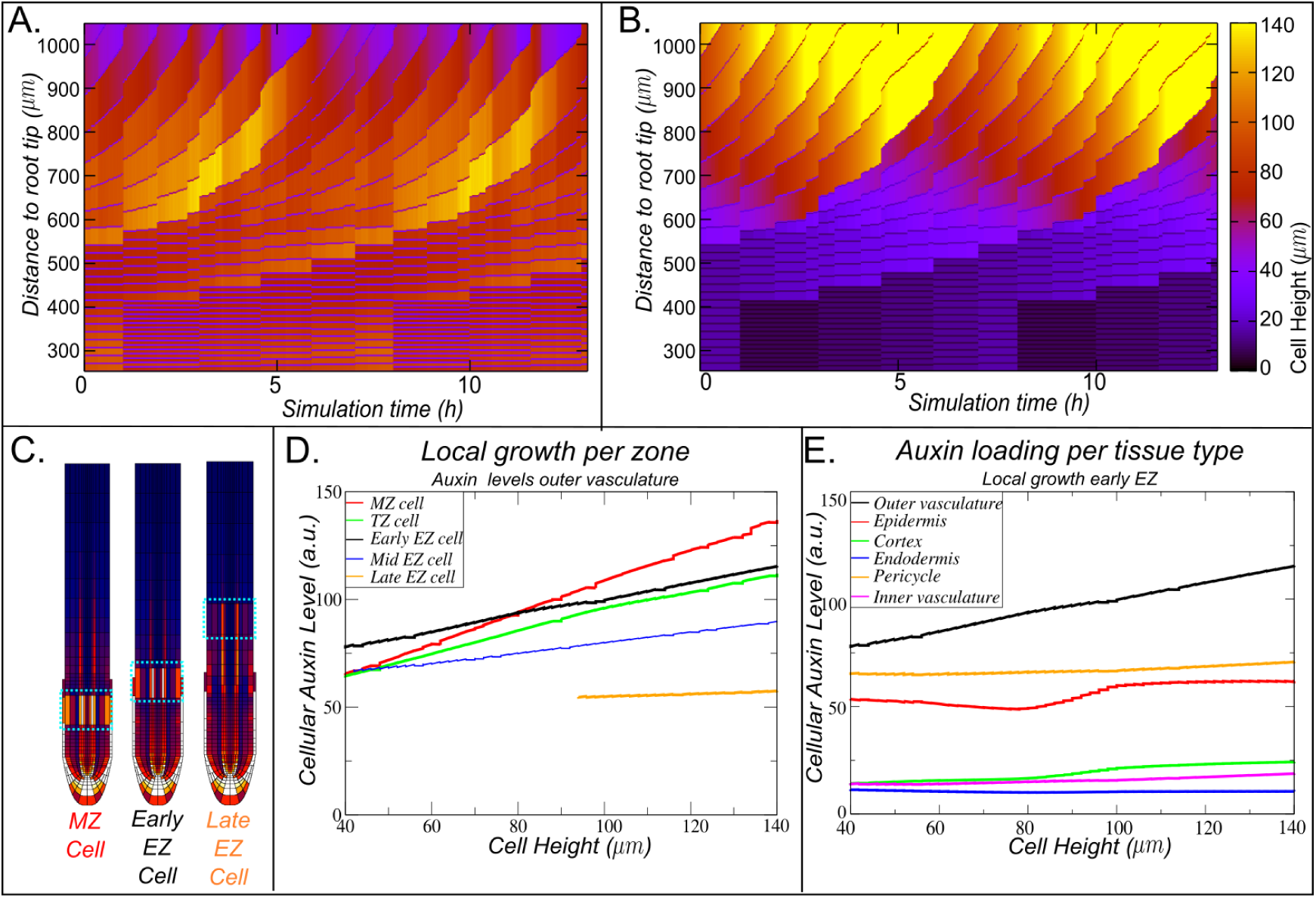
Cell size increases drive auxin increases. **A, B**. Auxin maximum coincides with larger sized cells arriving in EZ (compare A and B). **C**. Local growth simulations were performed to test the auxin loading capacity of growing cells in different zones, MZ, TZ (not shown), early EZ, mid EZ (not shown) and late EZ. **D**. Auxin concentration increases with cell size in all zones, magnitude of the increase depends on auxin availability being highest in MZ and lowest in late EZ and intermediate in TZ and early EZ, were highest availability occurs in the auxin loading zone at the end of the LRC. **E**. Auxin levels in different tissue for local growth in early EZ, outer vasculature shows most increase and highest levels of auxin

Our results show that auxin increase in the growing cells strongly correlates with cell size, independent of the growth location. Still, the level of increase does depend on local auxin availability, being highest in the MZ and lowest in the late EZ (Fig. 6D). Importantly, *in planta* meristematic cells grow only slowly and to a limited extent before dividing and becoming small again, and the fast and extensive growth necessary to load substantial amounts of auxin is restricted to cells residing in the elongation zone. Thus, combining our results with where cells actually undergo expansive growth, auxin loading occurs strongest in the early, most rootward regions of the EZ where high auxin and high growth rates coincide. Focusing specifically on elongation occurring in the early EZ, we next plotted temporal auxin dynamics in the different tissues of the root. Our results indicate that auxin loading preferentially occurs in the outer vasculature file (Fig. 6E).

Next, we aimed to answer why auxin content increases so strongly with cell size and does so preferentially in vasculature cells. We reasoned that upon cell size increase, an increase in the ratio of auxin uptake versus auxin release occurs. Due to the apolar nature of both active and passive auxin import, auxin import increases with membrane surface area and hence cell size. In addition, PIN exporters have a predominantly polar localization, with PINs mostly occurring on the non-growing basal (PIN1) and apical (PIN2) membranes, thereby causing a relative decrease in auxin efflux for increasing cell sizes. These effects will be strongest for vasculature cells, where due to their narrowness expansion causes the largest increase in surface to volume ratios. To test this hypothesis we first reduced the width of vasculature cells even further, increasing the number of vasculature cell files to maintain overall root topology constant. We indeed find a further increase in the amplitude of auxin oscillations in the vasculature (maximum level of ~120 versus ~90, ~33% increase), supporting the importance of the narrowness of vasculature cells (Fig 7A).

**Figure 7:**
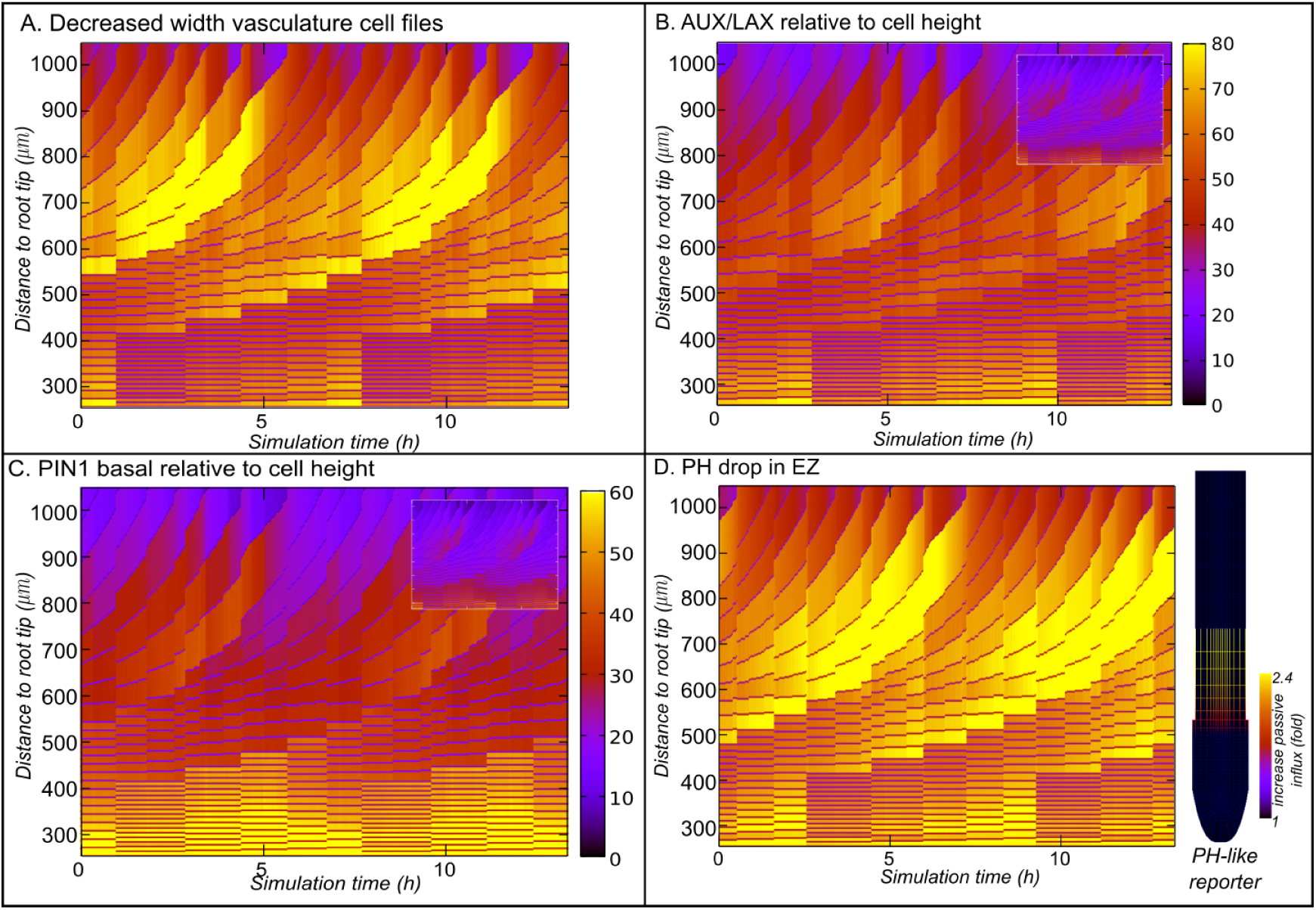
Vasculature cells have optimal loading capacity due to high surface to volume ratio.**A**. Width of vasculature cells was reduced with 2 μm, to keep root architecture 2 vasculature cell files were added. Amplitude of oscillations increase, indicating a positve correlation between cell width and auxin loading. **B**. AUX/LAX levels of EZ cells were normalized to cell height, effectively decreasing AUX/LAX levels up to 12 fold, a 22% reduction in amplitude is observed. **C**. PIN1 basal levels normalized to cell height, effectively increasing basal PIN1 levels levels up to 12 fold, a 33% reduction in amplitude is observed. **D**. PH dependent background auxin influx, PH-like reporter shows the relative increase in background influx compared to i_pas_. To compensate for the increased passive influx in the EZ, i_pas_ was reduced to 1.25. An increase of 33% in amplitude can be observed.

We then tested the importance of the different changes in auxin transport with cell size. In our default simulations, AUX/LAX membrane levels are assumed to stay equal during cell size increase, thereby effectively enhancing active uptake. However, it may be more reasonable to assume that differently sized cells express similar levels of AUX/LAX, resulting in a size dependent dilution of membrane levels. Therefore, we tested the effect of diluting AUX/LAX levels with cell height. Notably, in our model cells increase in height 12 fold while moving from the TZ towards and through the EZ, thus this normalisation substantially reduces AUX/LAX levels for large cells. Still, we find only a relatively minor reduction of auxin oscillation amplitude (maximum level of ~ 70 versus ~ 90, ~ 22% decrease), without qualitatively affecting priming dynamics (Fig 7B).

Following the same reasoning as for AUX1, cell-size independent PIN expression combined with non-growing apical and basal membranes on which these PINs are predominantly deposited would result in approximately constant apical or basal PIN levels. Thus, the cell size independence of apical and basal PIN levels in our model can be assumed to be realistic. We investigated the importance of the resulting decrease in effective efflux with cell size specifically for vascular cells. Therefore, we multiplied PIN1 levels with cell height, again resulting in changes of up to a factor of 12, to normalise efflux capacity with cell size. Under these conditions, a considerably more substantial decrease in oscillation amplitude was observed (maximum level of ~45 versus ~ 90, ~ 50% decrease) (Fig 7C).

Interestingly, it has been shown that in the TZ/EZ a significant drop in pH occurs, important for fast cell expansion. This pH drop causes a shift in the relative importance from AUX1 mediated active auxin import towards passive, background auxin influx [Rutschow et al., 2014, Street et al., 2016, Barbez et al., 2017]. Therefore, rather than normalizing background influx with cell size, we investigated the importance of background influx by incorporating a modest pH dependent increase of background influx with a factor of 2.4 in the elongation zone. Taking into account that this change in background auxin influx is 5 fold smaller in size than the cell height scaling induced changes for AUX/LAX and PIN1 we tested above, as well as the fact that even after this increase background influx is 3 fold less then active auxin uptake, this results in a significant increase in auxin oscillation amplitude (maximum level of ~125 versus ~90, ~39% increase) (Fig. 7D).

We conclude that auxin loading occurs preferentially in vasculature cells due to their narrowness, causing a more substantial increase in surface to volume ratios upon growth in these cells. The increased surface to volume ratio produces a dramatic increase in the auxin uptake capacity of these cells due to the increase of passive auxin uptake with membrane surface area and the relative decrease of polar PIN mediated efflux in larger cells. The possibly non-realistic increase in AUX/LAX-mediated transport with cell size present in our default model settings slightly contributed to but is not at all essential for this enhanced auxin uptake.

### 3.7 Cell size and loading time in the EZ determines auxin elevation

We thus found that the auxin reflux loop creates an auxin loading zone in the TZ and that long narrow cells have an advantage in terms of auxin loading potential. Inspecting the kymographs once more, we see that particularly large cells followed by a small cell have the largest auxin levels (Fig 6A and B). At the start of the EZ (270 um from the root tip), we see a periodic temporal sequence from small to large cells arriving at the EZ. Furthermore, we see that the cells that gain the highest auxin levels in Fig 6A are those cells that arrive with the largest size at the EZ, that reach their final size the earliest and that have the smallest cell following them (Fig 6a).

Theoretically, having a small cell below you may contribute to the auxin level of the above large cell in a total of three ways. First, nearby cells may compete for auxin, and given the lower auxin loading potential of a small cell, the above larger cells may be enabled to load more auxin. Second, due to exponential growth dynamics a smaller cell causes less displacement of its upper neighbour, allowing this cell to reside longer in the auxin loading region, we will refer to this as residence time. This larger residence time will allow the above cell to upload auxin for a larger time period, which may contribute to its overall auxin levels. Third, a larger residence time allows for a longer period of growth while inside this domain, so it will also allow the above cell to reach a larger size while in the auxin loading domain and may thus enhance auxin loading due to size. To disentangle these potential effects of competition, residence time and cell size we performed a series of artificially controlled growth simulations (see Methods section Local growth dynamics).

First, (Fig.8A, top), we varied elongation time while keeping all other settings constant, enabling us to investigate the impact of expansion rate and hence cell size attained within the TZ independent of changes in residence time. Next, (Fig 8A, middle), we varied meristematic cell cycle duration, thus impacting cumulative displacement rate and hence residence time in the EZ. To investigate the impact of residence time independent from the normally concomitant changes in cell size we applied a compensatory scaling of EZ elongation rate (for details see Methods section Local growth simulations). Finally, we varied whether the below cells divided or not. This allows us to investigate whether size dependent competition is relevant for auxin levels in the larger cells (Fig.8, bottom).

**Figure 8:**
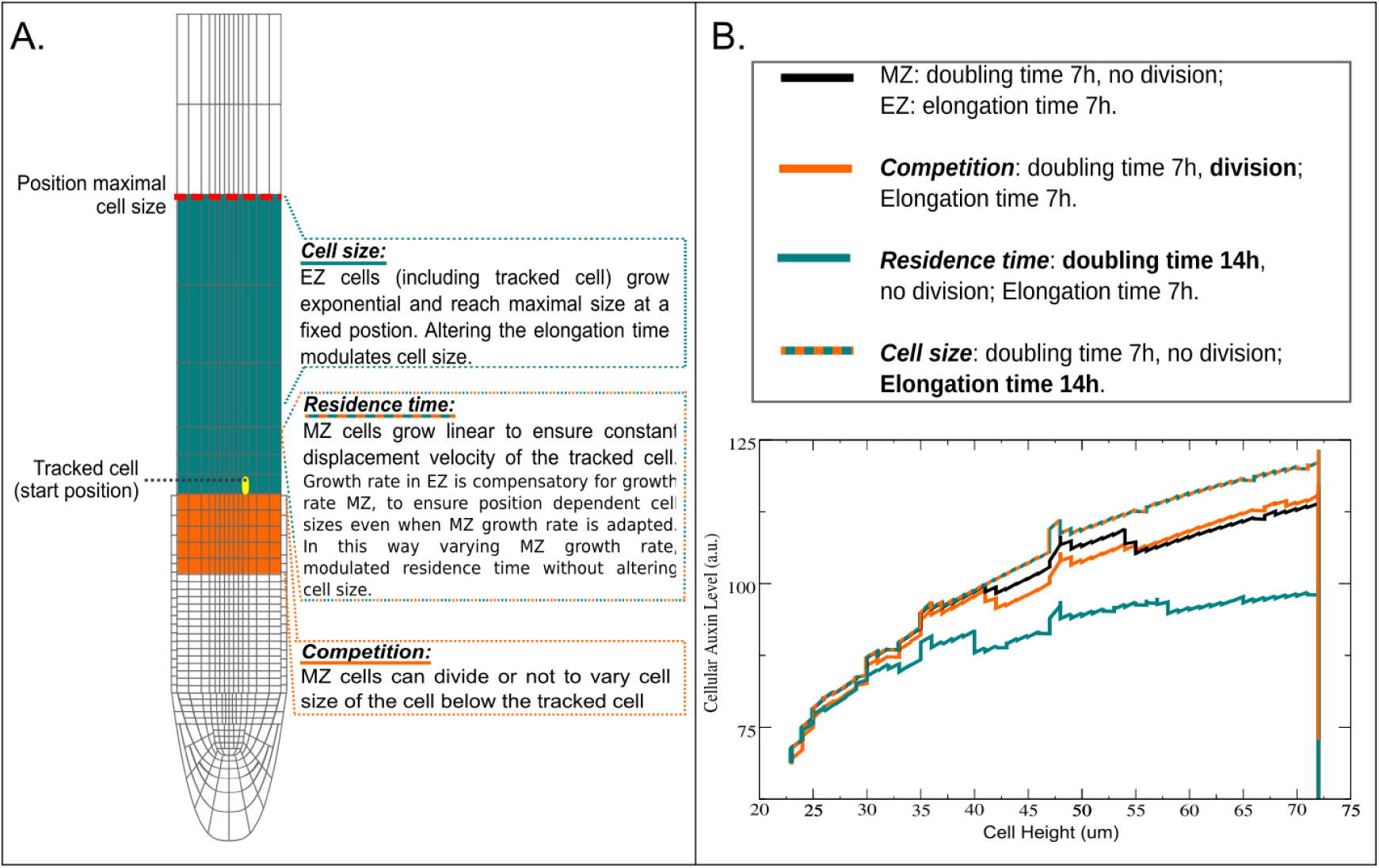
Artificially controlled growth experiment to disentangle potential effects of competition (Left bottem box), residence time (Left middle box) and cell size (Left upper box). Auxin level in the last cell of the MZ was tracked and a default simulation (Black line) was compared to the following variations: Cells in MZ below the tracked cell divided, enabling to investigate the influence of competition based on neighbor cell size (Orange line). Cells in the MZ grew slower, increasing the residence time of the tracked cell (Green Line). Elongation time was varied while keeping all other settings constant, enabling to investigate the impact of expansion rate and hence cell size attained within the TZ independent of changes in residence time (Green-Orange line). Indicating that size of neighboring cell hardly has an effect of auxin loading, while cell size has a large effect and residence time has a modest effect on auxin levels of a cell.

The results show that auxin loading hardly depends on the size of the neigbouring below cell, suggesting that reduced competition for auxin does not play a significant role. A modest increase in auxin loading can be observed for increased residence time. Finally, we see a significant reduction in auxin loading for a more slowly expanding, and hence smaller cells. Thus, the impact of a smaller below cell arises predominantly from the above cell having more time to grow and hence reaching larger sizes and more auxin loading potential, and to a lesser extent from the above cell spending more time in the auxin loading domain (Fig. 8B).

### 3.8 Growth dynamics drive cell size variations

The cell size variations we observe in our model arise as a natural consequence of the implemented longitudinal root zonation dynamics. Divisions of the lowermost meristematic cell, which can be interpreted as stem cells (for details see Methods), result in the maintenance of these lowermost cells while also producing new transit amplifying cells. Through subsequent divisions these transit amplifying cells give rise to growing clones of sibling cells while at the same time being pushed shootward through the continuous generation of new cells below them. Since the transition from division to expansion is based on longitudinal zonation, more shootward cells of a group of simultaneously growing cells will enter the expansion region and stop dividing earlier than the more rootward cells which can undergo another round of division. As a consequence, a large size difference arises between the most rootward cell just crossing the boundary and hence no longer dividing and the most shootward cell still dividing once more and hence becoming small again before entering the TZ/EZ (Fig S6).

However, in our model we thus far assumed cellular growth and division to occur at a constant, uniform rate across the meristem, causing perfect synchrony between clonal sibling cells. Experimental data show that cell cycle duration varies stochastically between cells. Additionally, cell cycle duration is known to depend on auxin levels, thus the root tip auxin gradient will translate into a gradient of cell cycle duration. Both these factors will cause cells to not grow and divide perfectly synchronously, possibly perturbing the above described process of alternating large and small cells. In Fig 9 we investigated the impact of a positional gradient dependent cell cycle duration(Fig. 9A) and stochasticity in cell cycle duration (Fig. 9B) on priming dynamics. We observe that oscillatory elevations in auxin levels remain, but occur with a somewhat reduced regularity in periodicity and amplitude. Indeed, priming oscillations only vanish when completely randomized growth and division dynamics were applied (Fig S7B).

**Figure 9:**
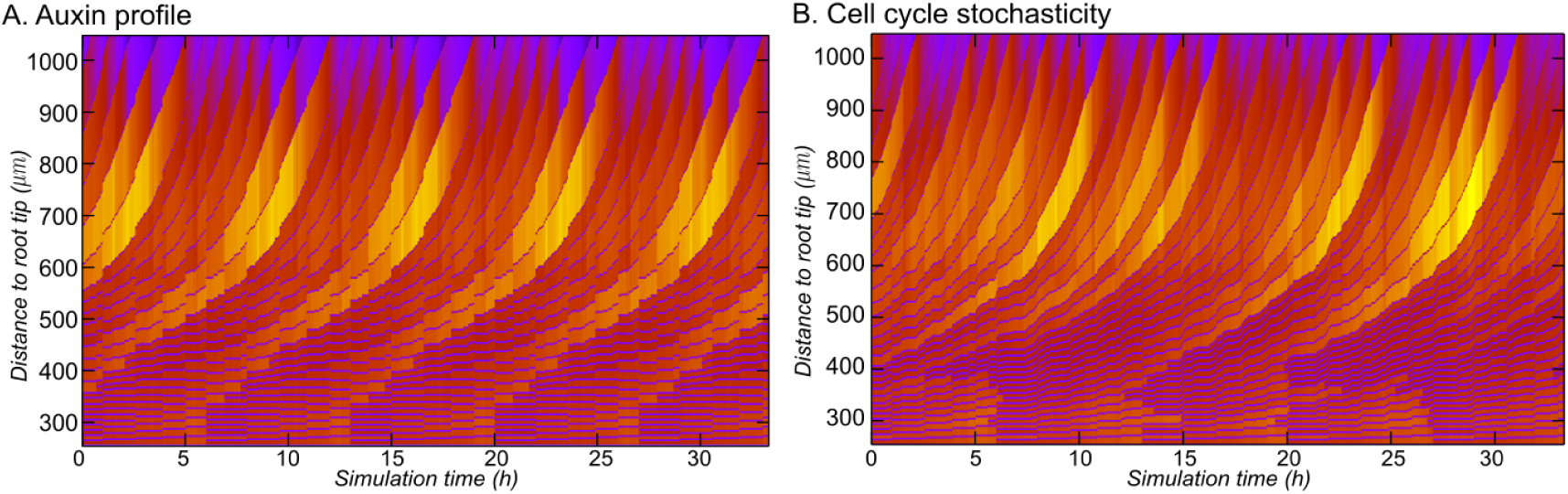
Cell cycle homogeneity is not a precondition for priming. **A**. Root with a positional gradient dependent cell cycle, emulating am auxin profile. **B**. Root with 10% stochasticity on cell cycle time.

Thus periodic priming and the underlying periodic variation in cell sizes arriving at the TZ requires sufficiently similar, but not identical, growth rates within groups of cells. Consistent with this, a recent experimental study tracking in detail cellular patterning during root tip growth uncovered similar semi-regular alternations in cell sizes at the start of the TZ (vonWangenheim2017) as we observe in our model when adding noise and/or position dependence on cell cycle duration.

### 3.9 Cell cycle dynamics determine priming frequency

In the previous section we explained how root tip growth dynamics induce periodic cell size variations. The largest cells arriving at the TZ arise from having had the longest time to grow after the previous round of division and just in time entering the TZ to not undergo a further round of division, whereas the directly following cells do undergo this extra division and hence arrive at their smallest. Thus, the frequency with which such pairs of successive large and small cells are generated is determined by the division frequency. Indeed, in the model we see that priming frequency is (approximately) equal to the 7h cell cycle duration, even if stochasticity or a modest distance dependent profile is added to modulate individual cell cycle durations (Fig 2, 9A and B). To further confirm this, we varied cell cycle durations, using values of 3, 5 and 9 hours, and in all cases priming period corresponded to cell cycle duration (Fig. S8)

To be more precise, priming depends on divisions *near the TZ boundary.* For approximately homogeneous division rates, the frequency of divisions near the TZ equals the overall cell cycle frequency in the MZ, and all of these divisions will result in a large cell followed by a small cell. Thus, priming frequency equals cell cycle frequency, as reported above. However, in case cell cycle durations differ significantly between stem cells and the distal meristem, the rate at which clones of sibling cells are generated differs from the rate at which these clones subsequently divide. This results in out of phase divisions of clones of cells that consecutively arrive at the TZ boundary. We hypothesize that this may affect priming frequency.

To investigate this we maintained cell cycle duration of most MZ cells at 7hrs, while varying (increasing) the cell cycle duration of only the lowermost, stem cell like, MZ cells. As an example, in figure S9C, for a stem cell cycle time of 12.25 hours, we observe a 12.25 hrs periodic pattern consisting of 2 nearby spaced peaks followed by a single further away peak, overall resulting in an average priming frequency of 6.125hrs. In contrast, for a stem cell cycle time of 9.45 hrs, a periodic pattern arises with 9.75 hrs between clear auxin peaks (Fig. S9B), but with a substantially lower level, additional auxin peak in between that is unlikely to result in priming. Thus, while the out of phaseness of divisions in consecutive clones causes the interval between divisions occurring close tot the TZ boundary to be lower then 7 hours this may cause either an increase or decrease in effective priming frequency. If cell cycle differences are sufficiently large, all division events have sufficient time to generate the necessary cell size differences and overall priming frequency will increase. If instead cell cycle differences are small, not all division events have time to result in sufficient cell size differences and effective overall priming frequency decreases.

Summarizing, cell cycle dynamics determine priming frequency. For significant differences between stem cell and bulk meristem cell cycles priming frequency can be both higher or lower than bulk meristem cell division frequency, while for approximately homogeneous division rates, priming frequency equals division frequency.

### 3.10 Meristem size influences LR spacing

In addition to cell division rates, the size of the meristem is an important determinant of root growth dynamics. For constant division rates, a larger meristem, with more growing and dividing cells, will result in a larger cumulative displacement velocity with which cells move towards and through the auxin loading zone. We hypothesize that this may also affect lateral root priming. First we investigate medium sized changes in meristem size, comparing priming dynamics for meristems of 25, 30 and 35 cells (Fig 10A, 2, 10B). Interestingly, we observe that while the temporal periodicity of auxin peaks remains constant at the 7h division frequency, and only minor changes in peak amplitude occur, the number of cells in between these auxin peaks clearly increases with meristem size. This can be understood as follows: while the auxin peaks arise from division events, the number of cells in between peaks is dependent on the number of cells that have passed by in the time interval between two division events. This number depends on cumulative displacement speed and thus depends on both meristem size and division rate.

**Figure 10:**
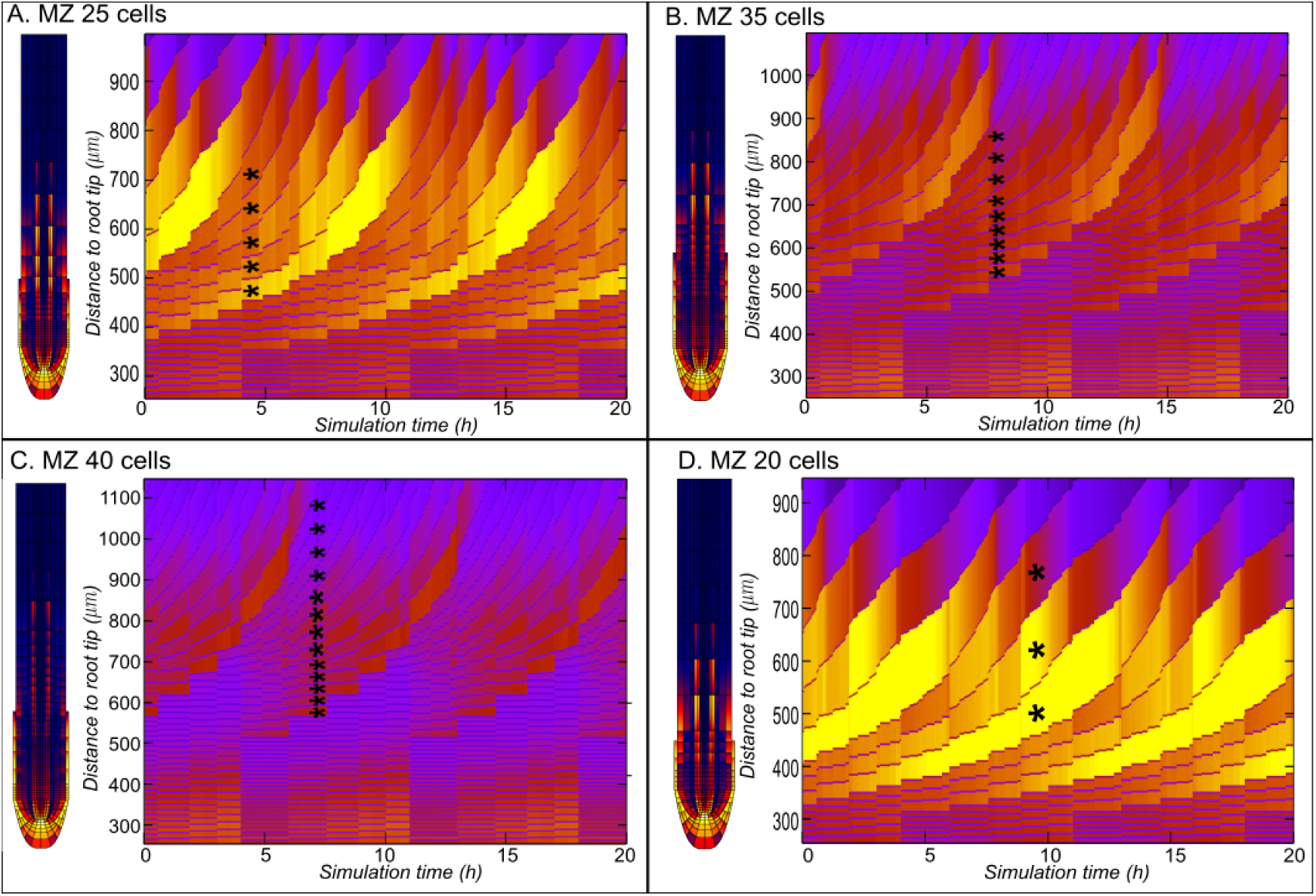
MZ size determines spacing of primed sites, but not the temporal frequency which is set by the cell cycle. All roots shown had a cell cycle of 7h, asterisks are to inidcate the number of cells passing per period. **A**. MZ with 25 cells, auxin amplitude but not frequency is increased compared to a default MZ of 30 (Fig. 2). Number of cells passing per period is reduced compared to default. **B**. MZ with 35 cells number of cell passing per period increases with the increased MZ size and auxin amplitude decreases. **C**. Extreme case of a large 40 cells MZ, number of cells passing per period is relative to the large MZ size. **D**. Extreme case of a small 20 cells MZ, number of cells passing per period is relative to the small MZ size.

Next, we investigated the impact of more extreme changes in meristem size, comparing the standard 30 cell meristem to meristems consisting of only 20 cells, and meristems of 40 cells. For very small meristems we see that almost all cells arriving in the auxin loading zone reach large cell sizes and load high amounts of auxin (Fig. 10C). Since small meristems result in low cumulative displacement rates, all cells have sufficient time to grow and load auxin in the TZ and thus hardly any periodicity in auxin levels arises. In contrast, for large meristems we see that almost all cells remain small and load little auxin (Fig. 10D), resulting from the limited growth time and auxin loading time caused by the high cumulative displacement speeds occurring in a large meristem. Again, hardly any periodicity in auxin levels occurs.

In summary, within a certain range of meristem sizes, our model predicts that meristem size affects the spacing of lateral root priming sites. Beyond that range, our model shows that if meristem sizes are too small or large and displacement speeds are too low or high, the initial cell size differences with which cells arrive at the TZ are insufficient to generate substantial cell size and auxin loading differences necessary for periodic priming.

## 4 Discussion

In this article we demonstrate a mechanism for LR priming that arises as an emergent property from root growth dynamics and root tip auxin reflux transport. We show that the auxin reflux loop generates an auxin loading zone at the basal meristem, with auxin loading occurring preferentially in the vasculature. For narrow vasculature cells cell expansion causes the most dramatic increase in surface to volume ratio. Due to the apolar nature of auxin uptake versus the polar localisation of auxin exporting PINs, this large increase in surface to volume ratio translates into a large increase in auxin import relative to export, causing large vasculature cells to have a very high auxin loading potential. In addition, vasculature cells tend to elongate more root-ward [Campbell and Turner, 2017, Rahni and Birnbaum, 2018], thus being longer than other cell types when arriving in the TZ. Passive auxin uptake in the TZ and early EZ is further enhanced by the steep decrease in pH accompanying elongation [Barbez et al., 2017, Street et al., 2016], which causes a shift in relative importance from active to passive auxin uptake [Rutschow et al., 2014].

For basal meristem auxin loading to result in priming oscillations, temporal variations in auxin loading caused by variations in cell sizes are critical. Using our model we demonstrate how these cell size variations arise naturally from root growth dynamics (Fig. S6). During root growth individual cells arise sequentially as progeny from the SCN [Bizet et al., 2015], and while moving through the MZ they give rise to expanding clones of sibling cells exhibiting approximately synchronous growth and division dynamics [von Wangenheim et al., 2017]. Due to the position based transition from the MZ to the TZ and EZ, these sibling cells sequentially leave the meristem. As a consequence the upper one leaves the MZ the earliest after the last division, and thus smallest in size. In contrast, the last sibling cell to enter the EZ before another round of division is up arrives the longest after the last division and is therefore largest in size. Either this sibling cell was the final cell of the clone, and the first cell of a new clone will follow next, starting again with a small cell that has recently undergone division. Alternatively, more sibling cells will follow but these will first undergo an extra round of division, also causing them to be small. Together, this leads to periodic differences in the size of cells when entering the TZ and EZ. Subsequently, these initial size differences become amplified due to the exponential growth dynamics in the EZ.

The LR priming mechanism we uncovered here predicts that priming frequency is determined by cell cycle duration. This prediction fits with the observation that in *cycd4;1* mutants, where specifically the pericycle cell cycle is slowed down, LR numbers are decreased [Nieuwland et al., 2009]. Furthermore, the observation that *aux1* mutants share with these mutants larger cell sizes and reduced LR numbers, suggests that in these mutants as well cell cycle length is increased [Nieuwland et al., 2009]. In addition, our results show that there is an important distinction between temporal spacing of priming sites, dependent on cell division dynamics, and spatial spacing of priming sites, which additionally depends on meristem size (Fig S8 and 10). In most studies, root architecture is typically described using density of LRs per root segment. Without the additional measurement of priming frequency and meristem size, this does not give insight in the source of changes in LR density. Indeed, we speculate that in many mutants and conditions in which a reduction of main root size and an increase in LR density is observed, for example phosphate starvation [Perez-Torres et al., 2008] or phloem differentiation mutants [Cattaneo and Hardtke, 2017], the increased LR density is at least partially caused by a smaller MR meristem causing decreased spacing between priming cells. Interestingly, in an extensive study investigating the effect of a range of concentrations of different nutrients on root architecture, clear inverse correlations were reported between MR length and first order LR density as well as first order LR length and second order LR density [Gruber et al., 2013]. These results are consistent with nutrient concentrations affecting meristem size, thereby influencing root lengths and LR density in opposite directions. Indeed, simulations with varying mersitem size show that smaller meristems have a higher density of primed sites compared to larger meristems (Fig. S10). This suggests that rather than conditions or a mutation having paradoxical opposite effects on MR and LR growth dynamics, the overall effect on root architecture may at least partly arise from one and the same change on MR growth dynamics. Finally, our results demonstrate that expansion rate, auxin production in the LRC and root tip, and auxin importer and exporter levels all modulate the amplitude of priming oscillations. The first does so by impacting the auxin loading potential of cells, the others by influencing the amount of auxin available for loading.

The priming mechanism described here fits well with earlier experimental observations demonstrating the importance of auxin transport [De Smet et al., 2007, Laskowski et al., 2008, Marhavý et al., 2013, Marhavý et al., 2014, Marhavý et al., 2016] and of auxin signaling in the vasculature [Dubrovsky et al., 2008, De Smet et al., 2010, Goh et al., 2012] for LR priming, as well as with research demonstrating the impact of IBA production on the amplitude, and to a lesser extent the frequency of priming oscillation [Xuan et al., 2015, Xuan et al., 2016]. According to our model, blocking auxin transport will abolish the reflux loop and hence the existence of a proper auxin loading domain, while blocking auxin signaling will abolish events downstream of auxin loading. Instead, blocking IBA production will mostly affect the amount of auxin available for loading, while on the long term it will also influence cell cycle dynamics, explaining as well the observed milder differences in priming frequency. The emergent nature of the proposed priming mechanism, combined with the experimentally observed partial redundancy of transporters [Vieten et al., 2005], also explains the limited effect single mutants in auxin transporters, such as *aux1* and *pin2*, have on lateral root numbers [De Smet et al., 2007, Swarup et al., 2008, Lewis et al., 2011, Xuan et al., 2015, Xuan et al., 2016]. Rather than fully abolishing auxin reflux and hence auxin loading, these single mutants merely affect reflux loop efficiency and hence the success rate of priming events.

How do our results fit with earlier proposed mechanisms of LR priming? Recently, Xuan et al (2016) proposed that priming dynamics are caused by the repetitive apoptosis of LRC cells, suggesting that these cells may export their auxin to the main root prior to their apoptosis. They reported a strong correlation between LRC apoptosis dynamics and priming dynamics. However, LR priming is only reduced in frequency and regularity in the apoptosis defective *smb* mutant. Based on our results we are tempted to speculate that the observed correlation of priming with LRC apoptosis arose from the general coordination of growth dynamics in the different tissues of the root tip. Earlier, Moreno-Risueno et al. (2010) reported oscillations in the dynamics of a large number of genes correlating with LR priming. Based on the intriguing parallels with vertebrate somitogenesis it was as the time proposed that a similar, cell-autonomous gene regulatory oscillator circuit could underly these oscillations [Moreno-risueno et al., 2010]. Thus far, no such oscillator circuit has been identified. Based on the results obtained here we propose that gene expression oscillations arise due to rather than being the cause of priming oscillations. Indeed, expression of many genes is either directly or indirectly auxin dependent [De Smet et al., 2007, Dubrovsky et al., 2008]. Importantly, this does not diminish the importance of these gene expression dynamics. As can be clearly seen in our simulations, reflux loop and growth dynamics together produce potent periodic auxin elevations but higher up towards the DZ auxin transport dynamics cause cells to loose these high auxin levels again. Thus, additional processes are required for the formation of stable prebranch sites with persistent high auxin signalling levels. We recently argued for the importance of distinguishing priming signal versus memory formation [Laskowski and ten Tusscher, 2017]. Based on our current results we suggest that the here described reflux and growth dynamics dependent auxin elevations constitute the priming signal, and that subsequent gene expression dynamics are critical for the memory formation that enables cells to maintain a high auxin response.

In summary, using a computational approach we demonstrated that oscillating auxin levels observed during LR priming are an emergent property of the auxin reflux loop architecture and root growth dynamics. Our model predicts that temporal priming frequency predominantly depends on cell cycle duration, while cell cycle duration together with meristem size control LR spacing. This prediction has important implications for research aimed at understanding root system plasticity. While traditionally, only LR density and root size is measured our results indicate the necessity of measuring cell cycle duration to be able to distinguish between variations in LR density arising from differences in cell cycle durations, meristem size or both.

## 5 Author Contributions

T.v.d.B. expanded the computer model, performed the simulation experiments, analysed the results and wrote the manuscript.

K.t.T. developed the computer model, conceived the study, analysed results and wrote the manuscript.

## 6 Acknowledgements

We would like to thank Ben Scheres, Viola Willemsen, Kavya Yalamanchili, and Marta Laskowski for helpful discussions and suggestions.

## References

Barbez, E., Dünser, K., Gaidora, A., Lendl, T., and Busch, W. (2017). Auxin steers root cell expansion via apoplastic pH regulation in Arabidopsis thaliana. Proceedings of the National Academy of Sciences, 114(24):E4884–E4893.

Beemster, G. T. and Baskin, T. I. (1998). Analysis of Cell Division and Elongation Underlying the Developmental Acceleration of Root Growth in *Arabidopsis thaliana*. Plant Physiology, 116(4):1515–1526.

Bennett, M. J., Marchant, A., Green, H. G., May, S. T., Sally, P., Millner, P. A., Walker, A. R., Schulz, B., Feldmann, K. A., Bennett, M. J., Marchant, A., Green, H. G., May, S. T., Ward, S. P., Millner, P. A., Walker, A. R., Schulz, B., and Feldmann, K. A. (1996). Arabidopsis AUX1 Gene: A Permease-Like Regulator of Root Gravitropism. American Association for the Advancement of Science, 273(5277):948–950.

Bhalerao, R. P., Eklöf, J., Ljung, K., Marchant, A., Bennett, M., and Sandberg, G. (2002). Shoot-derived auxin is essential for early lateral root emergence in Arabidopsis seedlings. Plant Journal, 29(3):325–332.

Bielach, A. and Benkova, E. Lateral root organogenesis — from cell to organ.

Bizet, F., Hummel, I., and Bogeat-Triboulot, M. B. (2015). Length and activity of the root apical meristem revealed in vivo by infrared imaging. Journal of Experimental Botany, 66(5):1387–1395.

Campbell, L. and Turner, S. (2017). Regulation of vascular cell division. Journal of Experimental Botany, 68(1):27–43.

Campilho, A., Garcia, B., Toorn, H. V., Wijk, H. V., Campilho, A., and Scheres, B. (2006). Time-lapse analysis of stem-cell divisions in the *Arabidopsis thaliana* root meristem. Plant Journal, 48(4):619–627.

Cattaneo, P. and Hardtke, C. S. (2017). BIG BROTHER Uncouples Cell Proliferation from Elongation in the Arabidopsis Primary Root. Plant and Cell Physiology, 58(9):1519–1527.

De Rybel, B., Vassileva, V., Parizot, B., Demeulenaere, M., Grunewald, W., Audenaert, D., Van Campenhout, J., Overvoorde, P., Jansen, L., Vanneste, S., Möller, B., Wilson, M., Holman, T., Van Isterdael, G., Brunoud, G., Vuylsteke, M., Vernoux, T., De Veylder, L., Inzé, D., Weijers, D., Bennett, M. J., and Beeckman, T. (2010). A novel Aux/IAA28 signaling cascade activates GATA23-dependent specification of lateral root founder cell identity. Current Biology, 20(19):1697–1706.

De Smet, I., Lau, S., Voss, U., Vanneste, S., Benjamins, R., Rademacher, E. H., Schlereth, A., De Rybel, B., Vassileva, V., Grunewald, W., Naudts, M., Levesque, M. P., Ehrismann, J. S., Inzé, D., Luschnig, C., Benfey, P. N., Weijers, D., Van Montagu, M. C. E., Bennett, M. J., Jüurgens, G., and Beeckman, T. (2010). Bimodular auxin response controls organogenesis in Arabidopsis. Proceedings of the National Academy of Sciences of the United States of America, 107(6):2705–10.

De Smet, I., Tetsumura, T., De Rybel, B., Frey, N. F. d., Laplaze, L., Casimiro, I., Swarup, R., Naudts, M., Vanneste, S., Audenaert, D., Inze, D., Bennett, M. J., and Beeckman, T. (2007). Auxin-dependent regulation of lateral root positioning in the basal meristem of Arabidopsis. Development, 134(4):681–690.

Dello Ioio, R., Nakamura, K., Moubayindin, L., Perilli, S., Taniguchi, M., T. Morita, M., Aoyama, T., Costantino, P., and Sabatini, S. (2008). A genetic framework for genetic control of cell division and differentiation in the root meristem. Science, 22:1380–1384.

Di Mambro, R., De Ruvo, M., Pacifici, E., Salvi, E., Sozzani, R., Benfey, P. N., Busch, W., Novak, O., Ljung, K., Di Paola, L., Marée, A. F. M., Costantino, P., Grieneisen, V. A., and Sabatini, S. (2017). Auxin minimum triggers the developmental switch from cell division to cell differentiation in the *Arabidopsis* root. Proceedings of the National Academy of Sciences, 114(36):E7641–E7649.

Dubrovsky, J. G., Gambetta, G. A., Hernández-Barrera, A., Shishkova, S., and González, I. (2006). Lateral root initiation in Arabidopsis: Developmental window, spatial patterning, density and predictability. Annals of Botany, 97(5):903–915.

Dubrovsky, J. G., Sauer, M., Napsucialy-Mendivil, S., Ivanchenko, M. G., Friml, J., Shishkova, S., Celenza, J., and Benková, E. (2008). Auxin acts as a local morphogenetic trigger to specify lateral root founder cells. Proceedings of the National Academy of Sciences of the United States of America, 105(25):8790–8794.

El-Showk, S., Help-Rinta-Rahko, H., Blomster, T., Siligato, R., Marée, A. F., Mähönen, A. P., and Grieneisen, V. A. (2015). Parsimonious Model of Vascular Patterning Links Transverse Hormone Fluxes to Lateral Root Initiation: Auxin Leads the Way, while Cytokinin Levels Out. PLoS Computational Biology, 11(10):1–40.

Eshel, A., Beeckman, T. (2013). Plant Roots: The Hidden Half, Fourth Edition. CRC Press, 4, illustr edition.

Goh, T., Kasahara, H., Mimura, T., Kamiya, Y., and Fukaki, H. (2012). Multiple AUX/IAA-ARF modules regulate lateral root formation: the role of Arabidopsis SHY2/IAA3-mediated auxin signalling. Philosophical Transactions of the Royal Society B: Biological Sciences, 367(1595):1461–1468.

Grieneisen, V. A., Xu, J., Maree, A. F. M., Hogeweg, P., and Scheres, B. (2007). Auxin transport is sufficient to generate a maximum and gradient guiding root growth. Nature, 449:1008–1013.

Gruber, B. D., Giehl, R. F. H., Friedel, S., and von Wiren, N. (2013). Plasticity of the Arabidopsis Root System under Nutrient Deficiencies. Plant Physiology, 163(1):161–179.

Herder, G. D., Van Isterdael, G., Beeckman, T., and De Smet, I. (2010). The roots of a new green revolution. Trends in Plant Science, 15(11):600–607.

Jensen, P. J., Hangarter, R. P., and Estelle, M. (1998). Auxin transport is required for hypocotyl elongation in light-grown but not dark-grown Arabidopsis. Plant Physiology, 116(2):455–462.

Kochian, L. V. (2016). Root architecture. Journal of Integrative Plant Biology, 58(3):190–192.

Kong, X., Zhang, M., De Smet, I., and Ding, Z. (2014). Designer crops: optimal root system architecture for nutrient acquisition. Trends in biotechnology, 32(12):597–8.

Laskowski, M., Biller, S., Stanley, K., Kajstura, T., and Prusty, R. (2006). Expression Profiling of Auxin-treated Arabidopsis Roots: Toward a Molecular Analysis of Lateral Root Emergence. Plant Cell Physiology, 47:788–792.

Laskowski, M., Grieneisen, V. A., Hofhuis, H., ten Hove, C. A., Hogeweg, P., Maree, A. F. M., and Scheres, B. (2008). Root system architecture from coupling cell shape to auxin transport.

Laskowski, M. and ten Tusscher, K. H. (2017). Periodic Lateral Root Priming: What Makes It Tick? The Plant Cell, 29(3):432–444.

Lewis, D. R., Negi, S., Sukumar, P., and Muday, G. K. (2011). Ethylene inhibits lateral root development, increases IAA transport and expression of PIN3 and PIN7 auxin efflux carriers. Development, 138(16):3485–3495.

Mahonen, A. P., Ten Tusscher, K., Siligato, R., Smetana, O., and Diaz-Trivino, S. (2014). PLETHORA gradient formation mechanism separates auxin responses. Nature, 0:1–5.

Malamy, J. E. and Benfey, P. N. (1997). Organization and cell differentiation in lateral roots of Arabidopsis thaliana. Development (Cambridge, England), 124(1):33–44.

Marhavý, P., Duclercq, J., Weller, B., Feraru, E., Bielach, A., Offringa, R., Friml, J., Schwechheimer, C., Murphy, A., and Benková, E. (2014). Cytokinin controls polarity of PIN1-dependent Auxin transport during lateral root organogenesis. Current Biology, 24(9):1031–1037.

Marhavý, P., Montesinos, J. C., Abuzeineh, A., Van Damme, D., Vermeer, J. E., Duclercq, J., Rakusová, H., Nováková, P., Friml, J., Geldner, N., and Benková, E. (2016). Targeted cell elimination reveals an auxin-guided biphasic mode of lateral root initiation. Genes and Development, 30(4):471–483.

Marhavý, P., Vanstraelen, M., De Rybel, B., Zhaojun, D., Bennett, M. J., Beeckman, T., and Benková, E. (2013). Auxin reflux between the endodermis and pericycle promotes lateral root initiation. EMBO Journal, 32(1):149–158.

Moreno-risueno, A. M. A., Norman, J. M. V., Moreno, A., Ahnert, S.E., Benfey, P. N., Moreno-risueno, M. A., Norman, J. M. V., Moreno, A., Zhang, J., Ahnert, S. E., and Benfey, P. N. (2016). Oscillating Gene Expression Determines Competence for Periodic Arabidopsis Root Branching. 329(5997):1306–1311.

Moreno-risueno, M. A., Norman, J. M. V., Moreno, A., Zhang, J., Ahnert, S. E., and Benfey, P. N. (2010). Oscillating gene expression determines competence for periodic Arabidopsis root branching. Science, 329(September):1306–1312.

Nieuwland, J., Maughan, S., Dewitte, W., Scofield, S., Sanz, L., and Murray, J. a. H. (2009). The D-type cyclin CYCD4;1 modulates lateral root density in Arabidopsis by affecting the basal meristem region. Proceedings of the National Academy of Sciences of the United States of America, 106(52):22528–33.

Novák, D., Kuchařová, A., Ovecka, M., Komis, G., and Šamaj, J. (2016). Developmental Nuclear Localization and Quantification of GFP-Tagged EB1c in Arabidopsis Root Using Light-Sheet Microscopy. Frontiers in Plant Science, 6(January):1–15.

Omelyanchuk, N. A., Kovrizhnykh, V. V., Oshchepkova, E. A., Pasternak, T., Palme, K., and Mironova, V. V. (2016). A detailed expression map of the PIN1 auxin transporter in Arabidopsis thaliana root. BMC Plant Biology, 16(1):1–12.

Perét, B., Swarup, K., Ferguson, A., Seth, M., Yang, Y., Dhondt, S., James, N., Casimiro, I., Perry, P., Syed, A., Yang, H., Reemmer, J., Venison, E., Howells, C., Perez-Amador, M. A., Yun, J., Alonso, J., Beemster, G. T. S., Laplaze, L., Murphy, A., Bennett, M. J., Nielsen, E., and Swarup, R. (2012). AUX/LAX Genes Encode a Family of Auxin Influx Transporters That Perform Distinct Functions during Arabidopsis Development. The Plant Cell, 24(7):2874–2885.

Perez-Torres, C.-A., Lopez-Bucio, J., Cruz-Ramirez, A., Ibarra-Laclette, E., Dharmasiri, S., Estelle, M., and Herrera-Estrella, L. (2008). Phosphate Availability Alters Lateral Root Development in Arabidopsis by Modulating Auxin Sensitivity via a Mechanism Involving the TIR1 Auxin Receptor. the Plant Cell Online, 20(12):3258–3272.

Rahni, R. and Birnbaum, K. D. (2018). Long-term time lapse imaging of Arabidopsis roots. bioRxiv.

Rogers, E. D. and Benfey, P. N. (2015). Regulation of plant root system architecture: Implications for crop advancement. Current Opinion in Biotechnology, 32(Figure 1):93–98.

Rutschow, H. L., Baskin, T. I., and Kramer, E. M. (2014). The carrier AUXIN RESISTANT (AUX1) dominates auxin flux into Arabidopsis protoplasts. New Phytologist, 204:536–544.

Strader, L. C. and Bartel, B. (2011). Transport and metabolism of the endogenous auxin precursor indole-3-butyric acid. Molecular Plant, 4(3):477–486.

Street, I. H., Mathews, D. E., Yamburkenko, M. V., Sorooshzadeh, A., John, R. T., Swarup, R., Bennett, M. J., Kieber, J. J., and Schaller, G. E. (2016). Cytokinin acts through the auxin influx carrier AUX1 to regulate cell elongation in the root. Development, 143(21):3982–3993.

Swarup, K., Benkova, E., Swarup, R., Casimiro, I., Peret, B., Yang, Y., Parry, G., Nielsen, E., De Smet, I., Vanneste, S., Levesque, M. P., Carrier, D., James, n., Calvo, V., Ljung, K., Kramer, E., Roberts, R., Graham, N., Marillonnet, S., and et al Patel, K. (2008). The auxin influx carrier LAX3 promotes lateral root emergence. Nature Cell Biology, 10:946–954.

Swarup, R., Friml, J., Marchant, A., Ljung, K., Sandberg, G., Palme, K., and Bennett, M. (2001). Localization of the auxin permease AUX1 suggests two functionally distinct hormone pathways operate in the Arabidopsis root apex. Genes Development, 15:2648–2653.

Swarup, R., Kramer, E. M., Perry, P., Knox, K., Leyser, O., Beemster, G. T., Haseloff, J., Bhalerao, R., and Bennett, M. J. (2005). Root gravitropism requires lateral root cap and epidermal cells for transport and response to a mobile auxin signal. Nature Cell Biology, 7(11):1057–1065.

Van Den Berg, T., Korver, R. A., Testerink, C., and Ten Tusscher, K. H. W. J. (2016). Modeling halotropism: a key role for root tip architecture and reflux loop remodeling in redistributing auxin. Development, 143:3350–3362.

Vieten, A., Vanneste, S., Wisniewska, J., Benková, E., Benjamins, R., Beeckman, T., Luschnig, C., and Friml, J. (2005). Functional redundancy of PIN proteins is accompanied by auxin-dependent cross-regulation of PIN expression. Development (Cambridge, England), 132(20):4521–4531.

von Wangenheim, D., Hauschild, R., Fendrych, M., Barone, V., Benková, E., and Friml, J. (2017). Live tracking of moving samples in confocal microscopy for vertically grown roots. eLife, 6.

Willemsen, V., Bauch, M., Bennett, T., Campilho, A., Wolkenfelt, H., Xu, J., Haseloff, J., and Scheres, B. (2008). The NAC Domain Transcription Factors FEZ and SOMBRERO Control the Orientation of Cell Division Plane in Arabidopsis Root Stem Cells. Developmental Cell, 15(6):913–922.

Xuan, W., Audenaert, D., Parizot, B., Müller, B. K., Njo, M. F., De Rybel, B., De Rop, G., Van Isterdael, G., Mühünen, A. P., Vanneste, S., and Beeckman, T. (2015). Root cap-derived auxin pre-patterns the longitudinal axis of the arabidopsis root. Current Biology, 25(10):1381–1388.

Xuan, W., Band, L. R., Kumpf, R. P., Van Damme, D., Parizot, B., De Rop, G., Opdenacker, D., Möller, B. K., Skorzinski, N., Njo, M. F., De Rybel, B., Audenaert, D., Nowack, M. K., Vanneste, S., and Beeckman, T. (2016). Cyclic programmed cell death stimulates hormone signaling and root development in Arabidopsis. Science, 351(6271):384–387.

